# Non-cell-autonomous mechanisms of tumor initiation and relapse by chromosomal instability

**DOI:** 10.1101/2025.10.31.685838

**Authors:** Simona Lafirenze, Bastiaan van Gerwen, Ajit I. Quirindongo, Aniek Janssen, Banafsheh Etemad, Pim Toonen, Wilma H.M. Hoevenaar, Natasja Costermans, Sameh Youssef, Alain de Bruin, Lodewijk A. A. Brosens, Nannette Jelluma, Geert J.P.L. Kops

**Author notes:** Correspondence to: GJPLK. Equal contribution (Simona Lafirenze, Bastiaan van Gerwen). joint senior authors (Nannette Jelluma, Geert J.P.L. Kops).

## Abstract

Chromosomal instability (CIN) is a hallmark of cancer, and a primary cause of genetic heterogeneity in tumors. Depending on the degree of CIN and the affected tissue, CIN can promote or suppress tumor formation, and high CIN induction has been proposed as a therapeutic strategy. How CIN achieves these effects is unclear. Here we use a conditional mouse model of graded CIN in combination with longitudinal monitoring of DMBA/TPA-initiated skin tumors to show that low CIN increases the frequency of skin tumor initiation, while higher CIN accelerates onset and growth rates. Strikingly, gene recombination analysis of the tumors reveals that upon high CIN induction the fast-growing tumors originate from rare low CIN cells, suggesting a strong non-cell-autonomous effect of high CIN. Gene expression analysis and immunohistochemistry show that high CIN causes epidermal hyperplasia, immune evasion and a regenerative response that stimulates low CIN tumor growth beyond what is achieved by induction of low CIN alone. Such cell-extrinsic effects may be a common mechanism of tumor formation by CIN, as we observe it also in CIN-induced tumors of the intestine, breast and mesentery. When high CIN is induced in established skin tumors, mimicking CIN-based therapy, tumors regress but relapse quickly. Relapsed tumors, too, arose from rare low CIN cells. Our findings have implications for our understanding of the contributions of CIN to cancer initiation and progression and give caution to the rationale for CIN therapies.

**Significance statement:** Chromosomal instability (CIN) is a source of genetic diversity in cancer. Different degrees of CIN can be highly oncogenic or tumor suppressive, the latter being a rationale for CIN-based therapies. Using a mouse model for precise control of CIN levels, we show that low CIN promotes tumor initiation, while moderate CIN accelerates both initiation and tumor growth. Surprisingly, moderate CIN acts through its effects on the surrounding tissue rather than directly within tumor cells, by triggering a regenerative response accompanied by immune evasion. When very high CIN is induced in existing tumors, tumors initially regress but rapidly relapse. These findings inform on the contributions of CIN to cancer initiation and progression and give caution to the rationale for CIN therapies.

## Introduction

Aneuploidy is a hallmark of human tumors (1–3). Aneuploidy is caused by errors in chromosome segregation during cell division, the frequency of which is relatively high in human cancer cells (1, 4, 5). This phenotype, known as chromosomal instability (6), can drive tumor formation and progression, as well as have tumor suppressive effects (6–18). The mechanisms by which CIN promotes tumor formation and progression are not well understood. CIN results in chromosomal copy number alterations and can thereby promote gains of oncogenes or loss of tumor suppressor genes (19). CIN also creates micronuclei, which are a source of oncogenic genome rearrangements, gene amplification, and heritable epigenetic alterations to the micronucleated chromosomes (20–24). Besides these cell-intrinsic processes, CIN can cause non-cell-autonomous effects. Through generation of gene dosage imbalances and micronuclei, CIN can elicit innate immune responses that on the one hand promote immune surveillance and on the other hand drive IL6-dependent survival of CIN cells as well as immune evasion and metastatic cell behavior (11, 25–30). The pro- vs anti-tumor effects of CIN seem to depend on tissue context, degree of CIN, and duration of the CIN phenotype, all potentially influencing proliferation and survival as well as tumor microenvironmental and immune responses. The various consequences of CIN thus contribute to genome destabilization, intra-tumor heterogeneity, and tumor-immune interactions, which are thought to enable cancer cells to evolve new traits and adjust to their changing environment during growth, progression, and treatment (13, 26, 31–33).

Using a mouse model for CIN termed CiMKi (<u>C</u>re-inducible *<u>M</u>ps1* <u>K</u>nock-in), we recently showed that the oncogenic potential of CIN depends on its level and location: moderate CIN is potently oncogenic in both small intestine and colon, but high CIN was so only in the colon (34). Very high CIN can however also be detrimental to tumors (10, 35). This spurred the notion that CIN could be used as therapy, in line with the effect of mainstay spindle poison therapies (e.g. paclitaxel). Indeed, several approaches to increase CIN in tumor treatments, for example by compromising the spindle assembly checkpoint (36) or mitotic motor proteins, have been tested and are currently under investigation in clinical settings (reviewed in (37, 38) and see clinicaltrials.gov). Here we use our CiMKi model of conditional, graded CIN and a classic skin cancer model that enables longitudinal monitoring of growth dynamics to examine the mechanisms by which CIN contributes to initiation and growth of primary tumors as well its impact on pre-established tumors.

## Results

### Induction of various degrees of CIN in skin

Our CiMKi mouse model for graded CIN allows tissue-specific induction of various degrees of CIN by conditional expression of different activity mutants of the master regulator of the SAC, the kinase Mps1. The resulting mutant combinations cause very low (WT/TA), low (WT/KD), moderate (TA/TA), high (TA/KD) and very high (KD/KD) CIN (Figure 1a). To examine the contributions of CIN to tumor formation, we combined CiMKi with a classic carcinogen model for cutaneous squamous cell carcinoma (cSCC), which enables accurate and longitudinal measurements of onset and growth of tumors after timed induction of various degrees of CIN. We previously showed that diverse degrees of CIN can be induced with our CiMKi model in mouse embryonic fibroblasts, in organoid cultures derived from colon and small intestine, and in the intestinal tissues (34). To verify its applicability for skin, we cultured skin organoids from normal, untreated dorsal skin (39) from *CiMKi;Rosa26-CreER^T2^*mice (Fig. S1a), and introduced H2B-mNeon by lentivirus to enable live-imaging of chromosome segregation and quantification of CIN (Fig. S1b). Addition of 4-hydroxy-tamoxifen (4-OHT) to the organoids efficiently induced Cre-mediated recombination of the *CiMKi* alleles (Fig. S1c). Live imaging of chromosome segregation showed that induction of the various CiMKi genotypes resulted in the same degrees of CIN (from very low to very high) as previously seen in the intestine (Fig. 1a, b). Painting dorsal skin of *CiMKi;Rosa26-CreER^T2^* mice with 4-OHT allowed for highly penentrant *in vivo* recombination of the *CiMKi* alleles specifically in the skin, as evidenced by undetectable signal of the wild-type nucleotide in the mutant codon position (Fig. 1d) We therefore conclude that our CiMKi model enables efficient induction of a range of CIN levels in the skin.

**Figure 1.**
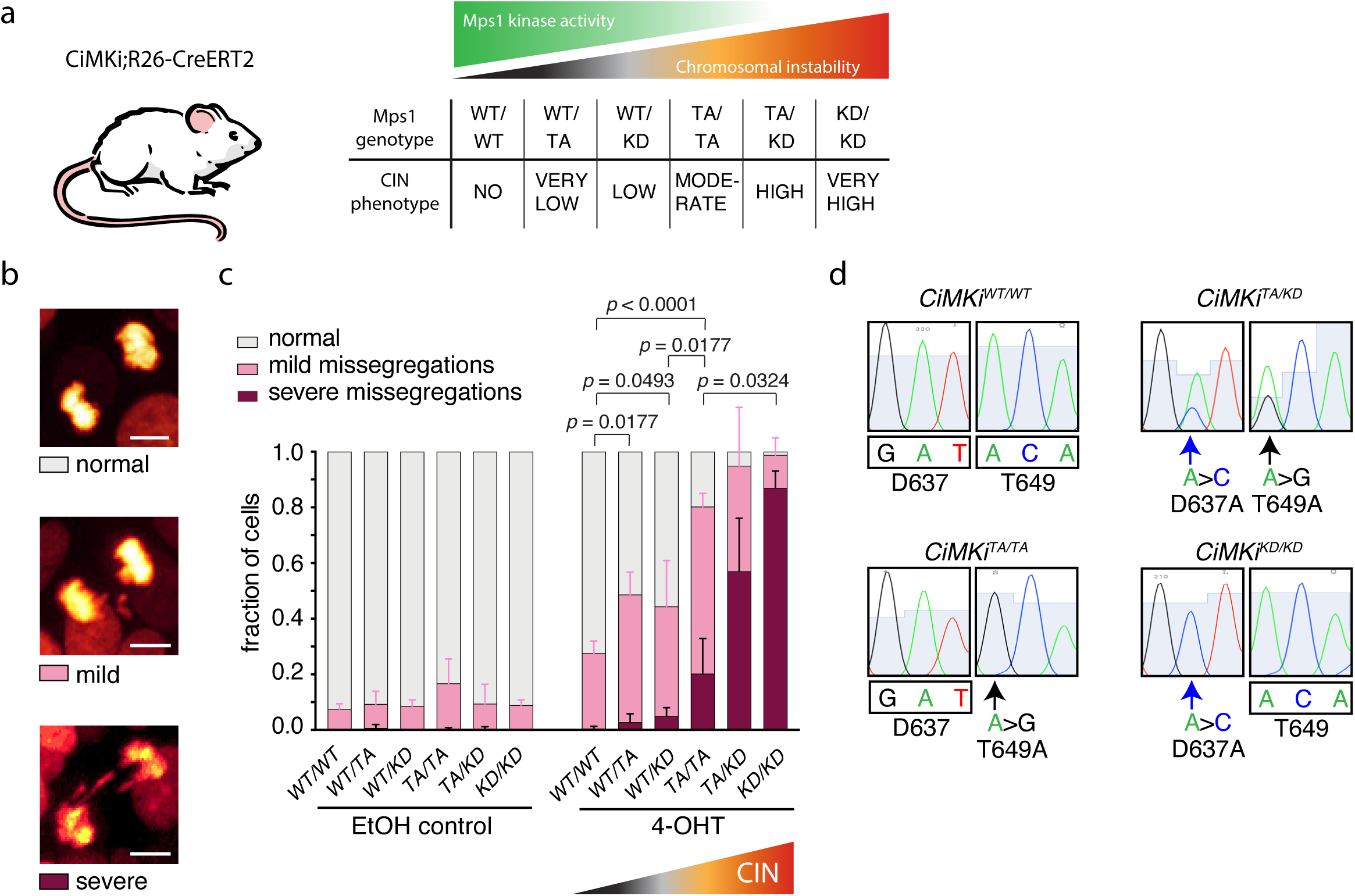
Induction of various degrees of CIN in skin. **a)** Schematic of the CiMKi mouse model. **b)** Stills from timelapse movies of skin organoids of the *CiMKi;Rosa26-CreER^T2^* genotypes 56 hours after 4-OHT addition. DNA is visualized by H2B-mNeon. Chromosome segregation errors were categorized as indicated: severe (≥three), mild (one or two). See also Supplementary Movies 1a, b. **c)** Quantification of chromosome segregation fidelity by time lapse imaging of skin organoids as in (4). Means and error bars ± SEM of three independent organoid lines per genotype (at least 25 divisions per line and at least three individual organoids per line). P-values are for differences between the totals of segregation errors (mild plus severe): ordinary one-way ANOVA (p<0.0001), followed by uncorrected Fisher’s LSD test, exact *p*-values are indicated when *p* < 0.05. **d)** Sequencing chromatograms of reverse transcribed RNA show efficient expression of both D637A (A to C) or T649A (A to G) CiMKi alleles in back skin of mice one week after topical treatment with 4-OHT.

### Different degrees of CIN have distinct effects on skin tumorigenesis

To examine the effects of various CIN levels on initiation and growth of skin tumors, we used the DMBA/TPA protocol for skin carcinogenesis on our *CiMKi;Rosa26-CreER^T2^* mice (40). In this protocol, a single application of the chemical initiator mutagen DMBA (7,12-dimethylbenz[*a*]anthracene) causes tumor initiation through oncogenic *Hras* (*41*), while subsequent repeated treatment with the phorbol ester TPA (12-O-tetradecanoylphorbol 13-acetate) then promotes clonal expansion of initiating cells to papilloma or squamous cell carcinoma (SCC). The exact mechanism of the TPA contribution is not well understood but likely involves activation of PKC and an overall stimulation of proliferation, survival and inflammation (42). We started biweekly application of TPA 7 days after DMBA treatment, followed by 5 single doses of 4-OHT on the backs of *CiMKi;Rosa26-CreER^T2^* mice with all CiMKi genotypes (Fig. 2a). This resulted in markedly different effects on various aspects of tumorigenesis. First, moderate and higher CIN levels, but not the lower CIN levels, caused substantially earlier onset of tumorigenesis (Fig. 2b, c, S2a). Second, these same CIN levels led to larger tumors, indicative of increased tumor growth rates (Fig. 2c, S2b). As a result, mice with moderate, high and very high CIN levels experienced higher average total tumor volume (Fig. 2c, S2c). End-point tumor grades were comparable between all CIN levels (Fig. S2d, e), but most mice from the moderate to very high CIN groups had to be sacrificed substantially earlier (Fig. 2b, S2f). Third, low and moderate degrees of CIN substantially increased the total number of tumors per mouse, whereas the higher degrees of CIN (high and very high) did so only very minimally (Fig. 2c). In summary: very low and low CIN levels affected only the frequency with which tumors formed (number of tumors), high and very high CIN accelerated tumor onset and growth rates (increased tumor size), and moderate CIN impacted all these features (numbers, size and onset).

**Figure 2.**
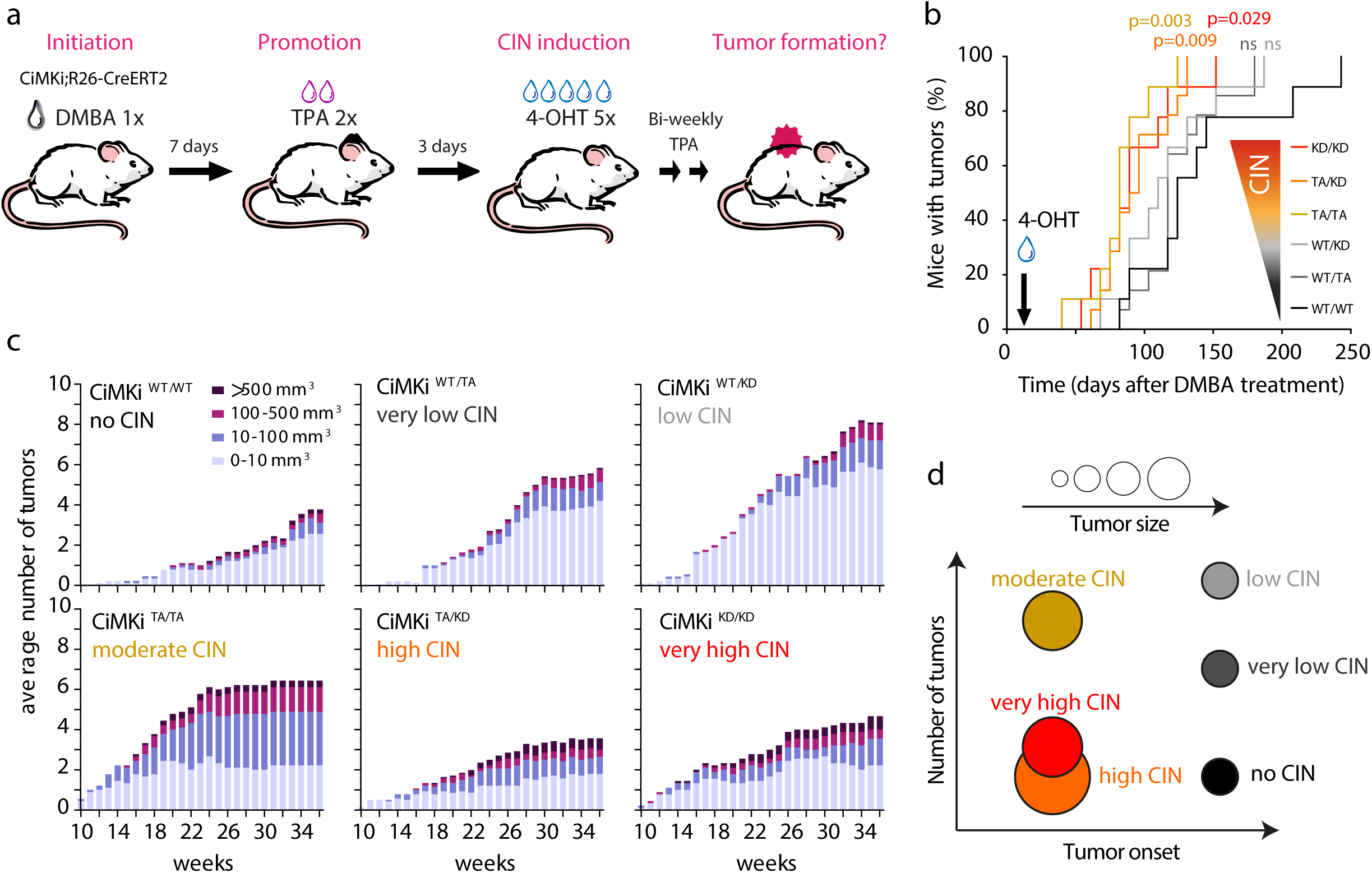
Different degrees of CIN have distinct effects on skin tumorigenesis. **a)** Treatment schedule of *CiMKi;Rosa26-CreER^T2^* mice. Mice were topically treated as indicated. **b)** Tumor incidence in *CiMKi;Rosa26-CreER^T2^* mice treated as in (a). n=9 (WT/WT, WT/KD, TA/TA), n=14 (WT/TA, TA/KD), or n=8 (KD/KD); p-values indicate significant differences of curves compared to WT/WT control curve (log-rank (Mantel-Cox) test). **c)** Quantification of the average number and size of tumors in time in the *CiMKi;Rosa26CreER^T2^* mice from b (4). Average tumor numbers were binned by size in four categories (0-10 mm^3^, 10-100 mm^3^, 100-500 mm^3^, and > 500 mm^3^) for each genotype. **d)** Schematic representation of the distinct effects of the various CIN levels on tumorigenesis as shown in (a-c).

Whole genome sequencing of three end-stage tumors from each CIN group showed distinct chromosomal copy number alterations specifically in tumors induced with moderate, high, and very high CIN. Most tumors from the WT group exhibited amplification of *Kras* and *Hras*-containing chromosomes 6 and 7 (43, 44) (Fig. S2g) probably due to the DMBA treatment. However, tumors from moderate, high and very high CIN mice had additional whole chromosome alterations (e.g. gain of chromosome 16) that were absent from the lower CIN tumors or the WT tumors (Fig. S2g). The different effects of various CIN degrees on tumor initiation and growth were therefore accompanied by distinct whole chromosome gains.

### Skin tumors from high CIN induction arise from low CIN cells

Our results show that various degrees of CIN can have distinct effects on tumor initiation and growth, including tumor-promoting effects of the highest CIN levels as a result of complete Mps1 inactivation (KD/KD). This was surprising, as we expected that very high CIN levels would be detrimental to cell viability. We therefore analyzed whether the tumor cells indeed carried fully recombined *CiMKi* alleles. For this, targeted Sanger sequencing for the *Mps1* point mutations was done on bulk cDNA obtained from mRNA from the tumors, and relative frequency of an allele was estimated based on relative peak intensities for the altered bases in the *Mps1* codons D367 and T649 (Fig. 3a, b). Although not a quantitative measure, this provided an estimate of allele presence in the population, which sufficed for our purposes. Analysis of tumors that arose in skin of WT/TA and WT/KD mice expressed ∼50% mutant Mps1 (Fig. S3a), as expected from the efficient recombination seen *in vivo* (see Fig. 1d, bottom panels). Surprisingly, however, only ∼50% of *Mps1* mRNA molecules in tumors from KD/KD mice carried the KD mutation (Fig. 3c), showing that these tumors contained cells with unrecombined *CiMKi* alleles. We envisioned three scenarios to explain this: (1) either the tumors consisted of a mix of fully recombined tumor cells with non-recombined cells (for example infiltrating immune cells); (2) the tumors consisted of cells in which only one allele had recombined; (3) a combination of the first and second scenario. Strongly supporting the second scenario is the observation that tumors from TA/KD mice all showed expression of the WT allele in combination with either the KD or the TA mutation but never both (Fig. 3c). Moreover, while recombination was initially very efficient for all genotypes when tested on healthy skin (no DMBA/TPA treatment) (Fig. 1d, 3d, S3b), over time expression of one of the mutant alleles was lost in the TA/TA, TA/KD and KD/KD mice (Fig. 3d and S3b). We therefore conclude that moderate, high and very high CIN cells are efficiently created in our model but are overtaken by rare cells in which only one allele had recombined and therefore had a low or very low CIN genotype (Fig 3e). Yet, as shown in figure 2, the growth characteristics of these tumors were very different from those in which only one allele had recombined from the start (e.g. WT/TA or WT/KD).

**Figure 3.**
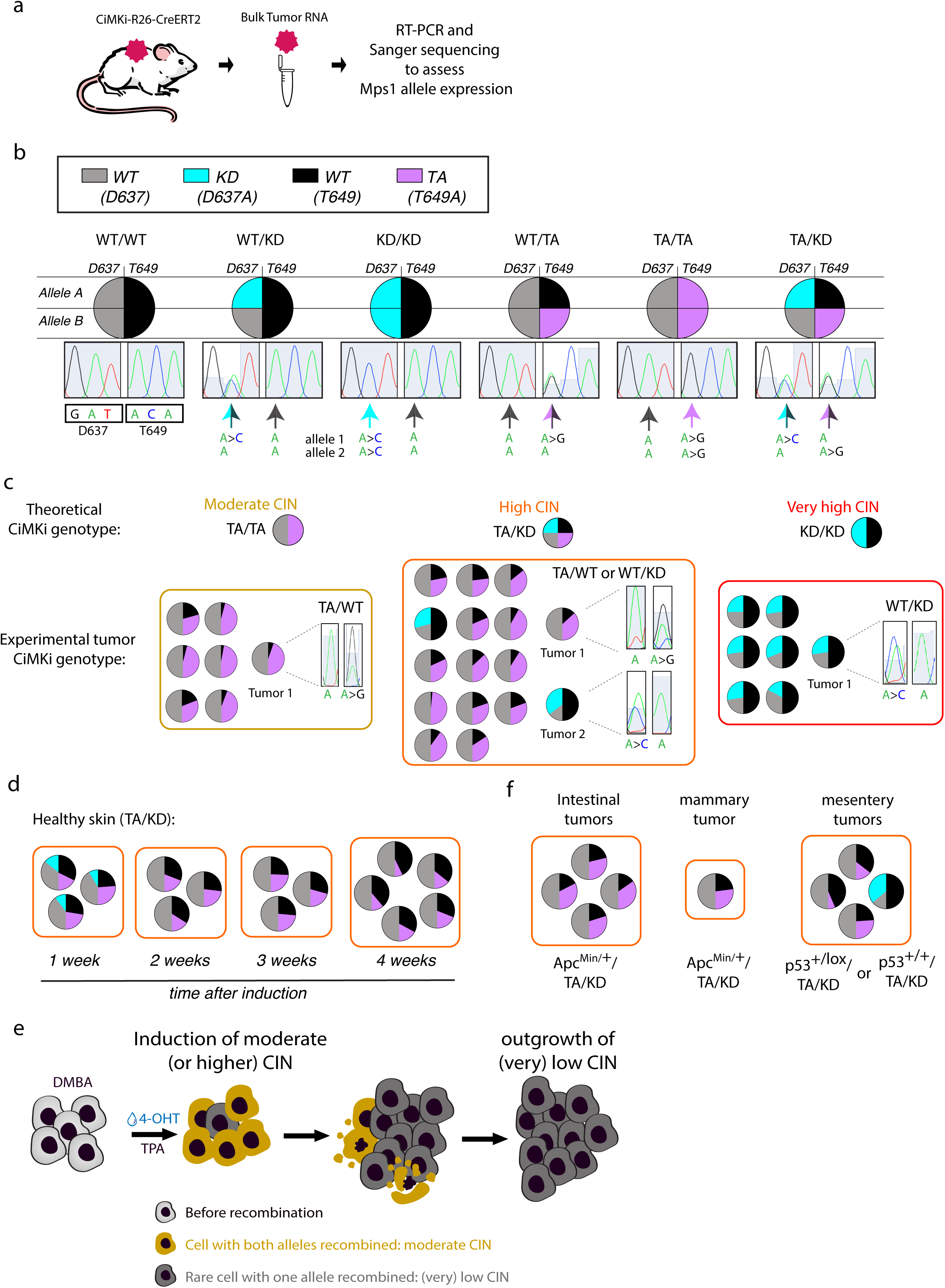
Skin tumors from high CIN induction consist of low CIN cells. **a)** Schematic representation of procedure: tumors are harvested from the back skin of the mice and RNA is extracted. RTPCR and Sanger sequencing follow to assess Mps1 allele expression. **b)** Schematic to indicate the theoretical percentage of mutant expression per CiMKi allele (A and B) for the different CiMKi genotypes, when recombination is complete. In the example for TA/KD, allele A is recombined to D637A (KD) and thus WT for the T649 site (blue/black), and allele B is recombined to T649A and thus WT for the D637 site (purple/gray). Sequencing chromatograms of cDNA show typical examples of sequences of the two mutation sites when fully recombined (arrows indicated mutation). **c)** Mutant expression in tumors from TA/TA (7 tumors), TA/KD (16 tumors) and KD/KD (7 tumors) mice. Notice the incomplete recombination of the Mps1 alleles in the tumors. **d)** Mutant expression in healthy skin from TA/KD mice, one, two, three, and four weeks after induction. See Fig. S3b for other *CiMKi* genotypes. **e)** Mutant expression in intestinal tumors and a mammary tumor from *Villin-Cre;* Apc^Min/+^; TA/KD mice, and from mesentery tumors from *Rosa26-CreER^T2^;* p53^flox/+^; TA/KD (1x) or *Rosa26-CreER^T2^;* p53^+/+^; TA/KD mice (3x). In b-d the same color code was used as in a). **f**) Moderate or higher CIN cells are efficiently created in CiMKi model after 4OH-T application, as detected by RTPCR and Sanger sequencing, indicating full recombination. However, in time these cells disappear and are overtaken by very rare, insufficiently recombined cells, in which only one allele had recombined and that therefore had a low or very low CIN genotype. These low CIN cells experience accelerated growth due to the microenvironment created by moderate/high CIN induction (Fig. 2d).

### Mammary, intestinal and mesentery tumors after high CIN induction consist of low CIN cells

Intrigued by the observation that high CIN greatly stimulates tumor forming capacity of rare low-CIN cells in the skin, we next wondered whether this also holds true for other tissues. To test this, we sequenced a variety of tumors from two studies in which high CIN also strongly promoted tumorigenesis: 1) intestinal and mammary tumors from *Apc^Min/+^; Villin-Cre; TA/KD* mice (34), and 2) mesentery tumors from *p53^lox/+^; Rosa26-CreERT2; TA/KD* or *p53^+/+^; Rosa26-CreERT2; TA/KD* (Fig. S3c,d). For these tumors, we observed similar *Mps1* mutant allele expression patterns as we did for the skin tumors (Fig. 3f), indicating that, again, the tumor-forming cells had only a single recombined *CiMKi* allele instead of two. This suggests that in various tissues, induction of high CIN accelerates outgrowth of rare, low CIN tumor cells.

### CIN can substitute for TPA in the skin carcinogenesis model

We next sought to better understand how high CIN can stimulate fast outgrowth of low CIN cells and reasoned it may be through mechanisms akin to the effects of TPA in promoting outgrowth of initiating cells (45). We therefore tested whether high CIN induction can replace TPA application in promoting tumorigenesis upon DMBA treatment. For this, *CiMKi;Rosa26-CreER^T2^* mice were treated with one initiating dose of DMBA, after which various CIN levels were induced (Fig. 4a) (46). While very low and low CIN did not impact tumorigenesis, induction of moderate and high CIN was sufficient to accelerate and enhance tumorigenesis by DMBA treatment (Fig. 4b,c): tumors grew earlier and bigger, although the numbers of tumors were not as high as when TPA was used (see upper left panel of fig. 2c). Very high CIN, while minimally affecting the number of tumors, caused tumors to grow larger and faster (Fig. 4b,c). CIN levels could therefore, to an extent, substitute for TPA as a tumor promoter.

**Figure 4.**
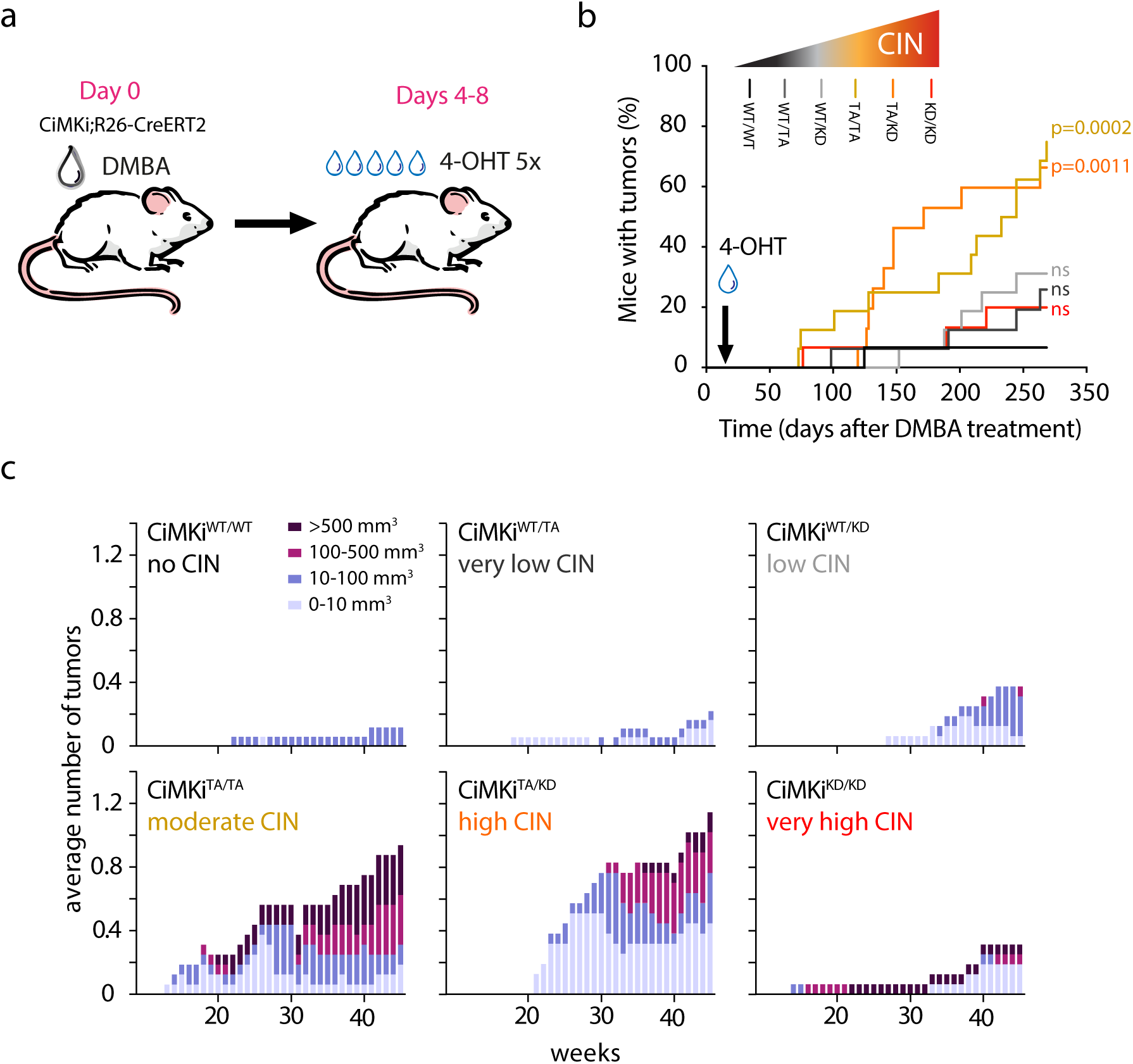
CIN can substitute for TPA in the skin carcinogenesis model. **a)** Treatment schedule of *CiMKi;Rosa26-CreER^T2^* mice. Mice were topically treated on their back skin with DMBA (one dose), then with 4-OHT once a day for five consecutive days. **b)** Tumor incidence in *CiMKi;Rosa26-CreER^T2^*mice treated as in (a). p-values indicate significant differences of curves compared to WT/WT control curve (log-rank (Mantel-Cox) test). **c)** Quantification of the average number and size of tumors in time in the *CiMKi;Rosa26CreER^T2^*mice from (b). Average tumor numbers were binned by size in four categories (0-10 mm^3^, 10-100 mm^3^, 100-500 mm^3^, and > 500 mm^3^) for each genotype.

### A minor role for inflammation in tumor promotion by CIN

Whereas TPA has been reported to induce a plethora of processes, a notable one is inflammation, a known risk factor for cancer progression (47–49). To directly test the contribution of inflammation to the tumor-promoting effects of high CIN, we repeated the experiment in which we used moderate and high CIN to replace TPA, but now while treating the mice throughout the experiment with the non-steroidal anti-inflammatory drug (NSAID) carprofen (Fig. 5a) (46). Immunohistochemistry confirmed that carprofen effectively inhibited CD45+ immune cell infiltration in TPA control and CIN-induced skin and skin tumors (Fig. 5b,d). As expected, carprofen reduced the ability of TPA to promote tumor promotion after DMBA application (Fig. 5b). Suprisingly, however, despite effective immune suppression, carprofen did not substantially reduce the number, size, or time of onset of tumors induced by moderate CIN, and only marginally of tumors induced by high CIN (Fig. 5c,d). We thus conclude that while inflammation may somewhat contribute to CIN-driven tumor promotion, it does not have a prominent role in our model.

**Figure 5.**
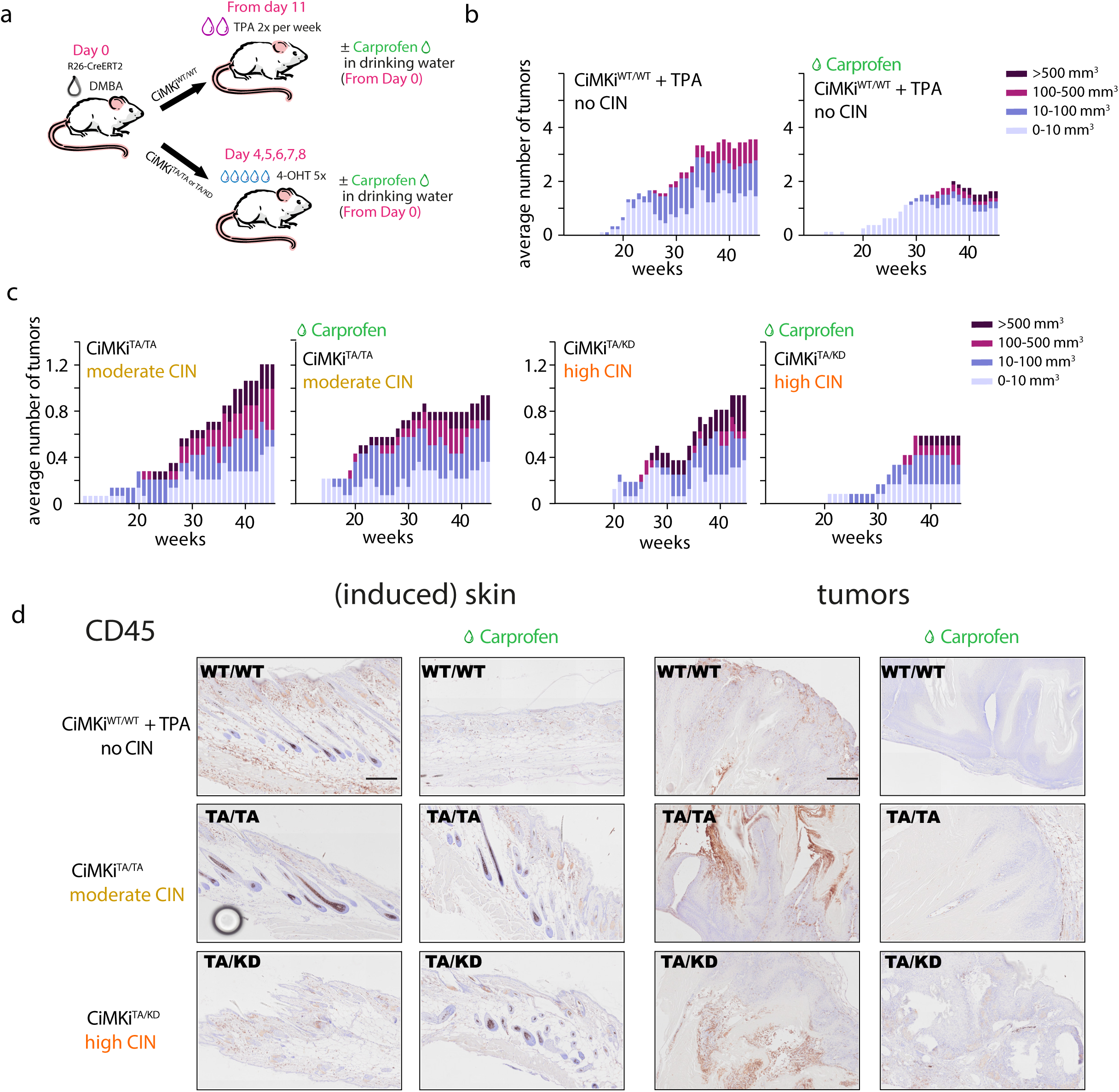
A minor role for inflammation in tumor promotion by CIN. **a)** Schematic representation of the experimental design: after DMBA application, no CIN (WT/WT) mice received biweekly treatment of TPA with or without carprofen in the water since the start of the experiment. These were taken along as a control for the effects of Carprofen. CIN (TA/TA and TA/KD) mice were treated with 4OH-T, after DMBA application, with or without carprofen in their drinking water. **b**) Quantification of the average number and size of tumors in time in no CIN (WT/WT) mice treated as in figure 4a, but with having carprofen in their drinking water for the duration of the experiment **c**) Quantification of the average number and size of tumors in time in *CiMKi;Rosa26CreER^T2^*moderate (TA/TA) and high (TA/KD) CIN mice treated as in figure 4a. Average tumor numbers were binned by size in four categories (0-10 mm^3^, 10-100 mm^3^, 100-500 mm^3^, and > 500 mm^3^) for each genotype. **d)** Decrease of immune (CD45+) cells in all carprofen treated conditions, both in induced skin and in tumors. Scale bar is 100 μm.

### Moderate/high CIN promotes a regenerative tissue response and epidermal hyperplasia

Our data imply that cells with high CIN, efficiently induced in our model, stimulate rare cells with low CIN to form tumors in a manner largely independent of inflammation. To identify the mechanism by which high CIN accomplishes this, we induced CIN in the skin for one and four weeks, without DMBA/TPA application, and subsequently performed bulk RNA sequencing (Fig. 6a, S4a). Analysis of differentially expressed genes (DEGs) (50) revealed that one week of CIN induction had caused an increase in keratinization and cellular proliferation, while fatty acid metabolism and cytokine expression were suppressed (Fig. 6b, S4b, c, d). A similar albeit more pronounced signature was seen four weeks after CIN induction (Fig. 6c, S4e, f).

**Figure 6.**
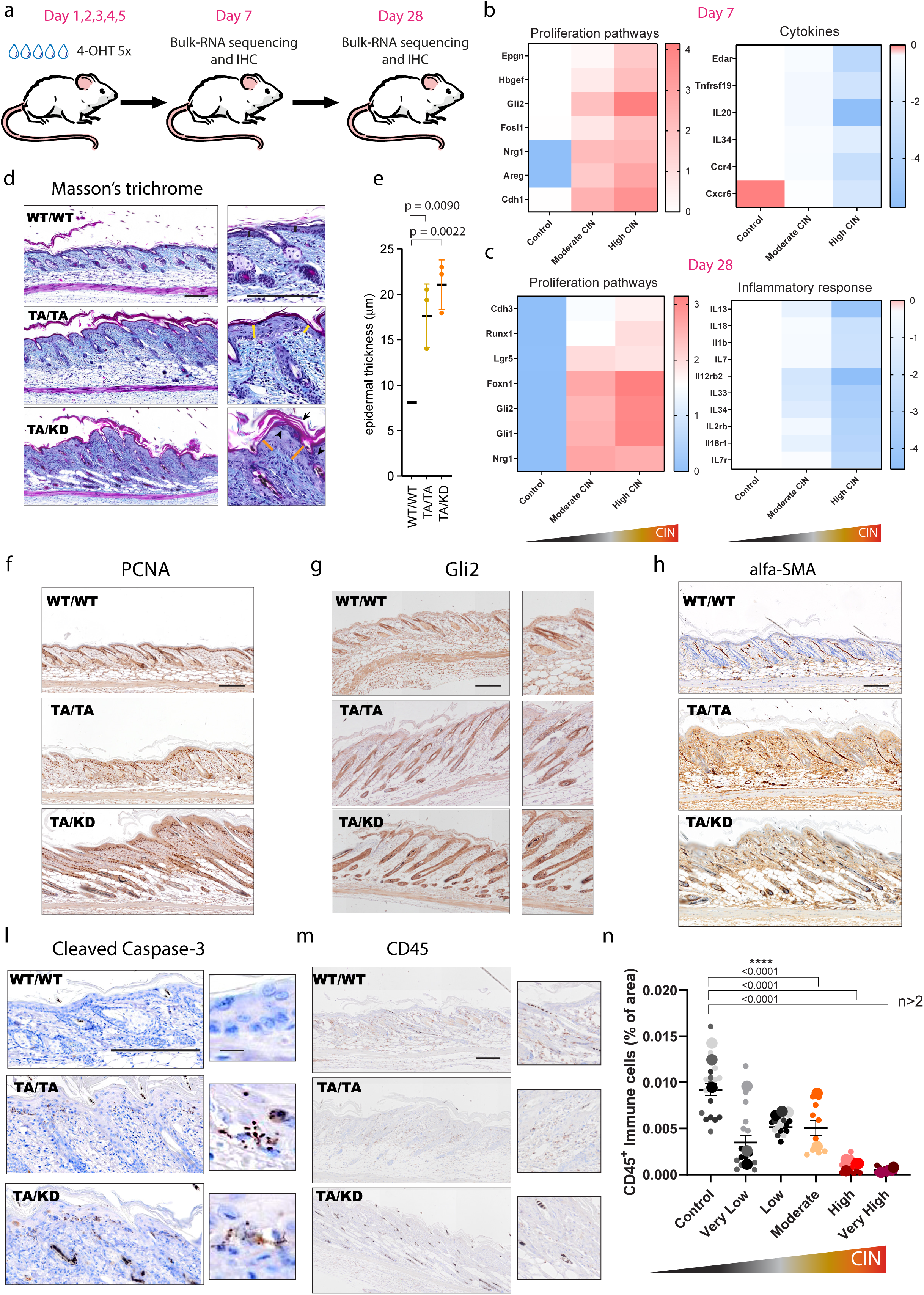
Moderate/high CIN promotes a regenerative tissue response and epidermal hyperplasia. **a)** Schematic representation of the experimental design: *CiMKi;Rosa26CreER^T2^* (no CIN, moderate and high CIN). Mice were topically treated on their back skin with 4-OHT once a day for five consecutive days. Skin tissues were harvested for bulk-RNA sequencing and for immunohistochemistry (IHC) at day 7 and at day 28 from CIN induction. **b)** Heatmap (red up, blue down) showing expression of upregulated proliferation genes and downregulated expression of cytokines in high CIN compared to no CIN skin at day 7 from CIN induction. **c**) Heatmap (red high, blue down) showing expression of upregulated proliferation genes and downregulated expression of inflammation in high CIN compared to no CIN skin at day 28 from CIN induction. **d)** Masson’s trichrome staining of mouse back skin one week after induction of moderate (TA/TA) or high (TA/KD) CIN, or without induction of CIN (WT/WT). In the TA/KD sample large amounts of keratohyalin granules (inset, arrowheads), and hyperkeratosis (inset, arrow) is seen. Also, enlarged cells and irregular nuclei, and epidermal hyperplasia (acanthosis, straight lines in insets indicate thickness of epidermis; see also (b)), are evident in both moderate and high CIN groups. In addition, we observed changes in the fibrotic structure (light blue staining) and signs of immune cell infiltration. **e)** Measurement of epidermal thickness shows significant hyperplasia in the moderate and high CIN groups, compared to the WT/WT control. Means and error bars ± SEM of three independent mice per genotype (20 measurements randomly distributed along epidermis per mouse). P-values are for means: ordinary one-way ANOVA (p<0.0001), followed by Tukey’s multiple comparisons test, exact *p*-values are indicated when *p* < 0.05. **f)** Increased proliferation in back skin one week after induction of moderate (TA/TA) or high (TA/KD) CIN, as judged by PCNA staining: in controls (WT/WT), positive cells are nicely structured in 1-2 layers of suprabasal cells along the epidermis and hair follicles, while CIN samples show disorganization and a more random distribution of positive cells within the dermis and more than two positive cell layers in the epidermis. **g)** Increased expression of Gli2 regenerative marker in all epithelial cells of the hair follicle (outer root sheath and dermal papilla) in moderate (TA/TA) and high (TA/KD) CIN samples compared to no CIN skin after 7 days from CIN induction. Scale bars are 100 μm **h)** Increased activated fibroblasts as judged by SMA staining in the back skin one week after induction of moderate (TA/TA) or high (TA/KD) CIN: Compared to controls (WT/WT), more SMA positive signal is observed, in dermis and/or epidermis. **i)** Increased apoptosis in the back skin one week after induction of moderate (TA/TA) or high (TA/KD) CIN as judged by cleaved caspase 3 (CC3). Punctate patterns of CC3 staining is observed mostly below the basal layer of the epidermis (insets), which are rarely observed in controls (WT/WT). **m**) Decrease of CD45+ cells in moderate (TA/TA) and high (TA/KD) CIN skin compared to no CIN after 7 days from CIN induction **n**) Quantification of CD45+ cells in *CiMKi;Rosa26-CreER^T2^* all ranges of CIN. Absence of CD45+ cells in all degrees of CIN, although to a bigger extent in high (TA/KD) and very high (KD/KD) CIN. Means and error bars ± SEM of three independent mice per genotype, except for moderate and very high which are two (6 regions of interest randomly distributed along epidermis per mouse). P-values are for all datapoints from all replicates: ordinary one-way ANOVA (p<0.0001), **(d, f-m)** Scale bars are 100 μm.

We next performed immunohistochemistry on skin tissues collected at the same timepoints. In line with the keratinization observed in the gene expression data, Masson’s trichrome staining showed significant epidermal hyperplasia (acanthosis) in both moderate and high CIN groups, as seen by thickening of the epidermis, enlarged cells and irregular nuclei, increased keratohyalin, and hyperkeratosis in all high CIN samples and in most moderate CIN samples (Fig. 6d,e,). All high CIN samples and the majority of moderate CIN samples exhibited hair follicle extension into adipose tissue, indicating anagen phase activation of hair growth, and this persisted after four weeks (Fig. 6d,f, S4d,e,g,h). Furthermore, we observed signs of tissue regeneration: cells positive for PCNA or ki67 (proliferation), Gli1 and Gli2 (skin development and regeneration) and SMA (fibroblast activation) were enriched in the suprabasal strata of the epidermis and partly in the hair follicle, exclusively in the moderate to very high CIN samples (Fig. 6f,g,h, S4i-n). These enrichments may reflect a response to increased apoptosis induced by high CIN, as cleaved caspase-3 was detected in the basal epidermal layer of moderate and high CIN samples but was rarely seen in the control (Fig. 6l, S4p). In line with suppression of cytokines in the gene expression analysis, we found depletion of CD45+ cells in the tissue, indicative of immune evasion (Fig. 6m,n, S4o). The anagen growth and the regenerative response, accompanied by the absence of immune cells, continued after four weeks from CIN induction (Fig S4e,g,h). These data suggest that induction of moderate to very high CIN levels (TA/TA, TA/KD, and KD/KD) in healthy skin triggers a proliferative and regenerative response that persist for weeks and leads to hyperplasia, which correlated with earlier tumor onset and enhanced tumor initiation and growth upon DMBA/TPA treatment (Fig. 2).

### Tumor relapse by low CIN cells after regression from high CIN induction

CIN induction is a therapeutic strategy for cancer (37, 51). Our data suggest that treatments that elevate CIN rates may have the unwanted consequence of promoting outgrowth of tumor cells with relatively low CIN. Since the DMBA/TPA model is well suited to monitor tumor growth longitudinally, we set out to test this by first establishing tumors by DMBA/TPA treatment of *CiMKi;Rosa26CreER^T2^* mice and then subsequently inducing moderate, high or very high CIN. When tumors reached a volume between 100-200 mm^3^, they were painted with 4-OHT to recombine the *CiMKi* alleles and induce CIN (Fig. 7a). All three CIN levels led to initial regression of the skin tumors, the extent positively correlating with the degree of CIN induced (Fig. 7b,c, S5a,b). Tumors relapsed after one to seven weeks after 4-OHT treatment (Fig. 7b, c), with the exception of two out of 19 tumors that were challenged with very high CIN (Fig. S5b, black lines in the KD/KD graph). Sequencing of *Mps1* cDNA again showed that relapsed tumors from KD/KD mice expressed ∼50% of the KD mutation, and 50% wildtype Mps1 (7 out of 7) (Fig. 7d). Similarly, 4 out of 5 tumors from TA/KD mice expressed either the KD (2 out of 5) or TA (2 out of 5) mutation in a near 1:1 ratio with wildtype *Mps1* (Fig. 7d), and we even observed two tumors from the same TA/KD mouse in which one expressed only T649A and the other one only D637A (Fig. S5c). Therefore, both during tumor promotion and tumor relapse, moderate, high and very high CIN accelerate outgrowth of tumor cells with low CIN.

**Figure 7.**
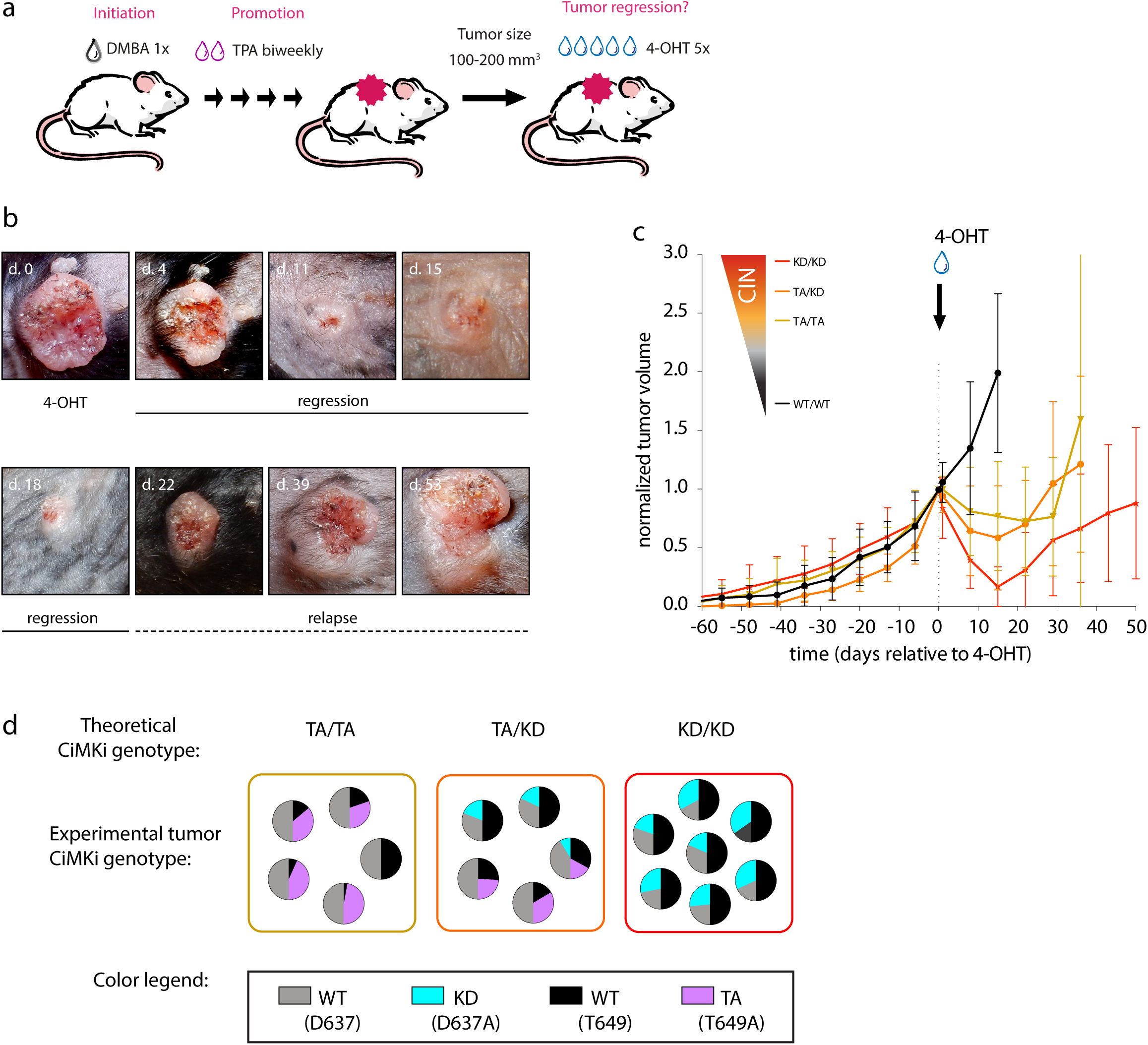
Tumor size dynamics after high CIN induction as a therapy mimic. **a)** Treatment schedule of *CiMKi; Rosa26-CreER^T2^* mice. Mice were topically treated on their back skin with DMBA (one dose), and TPA (biweekly) to promote tumor formation, until the end of the experiment. Tumors sized 100-200 mm^3^ were painted with 4-OHT on five consecutive days to induce the *CiMKi* alleles specifically in the tumors. **b)** Example of a regressing and relapsing tumor in a *CiMKi^KD/KD^*; *Rosa26-CreER^T2^* after 4-OHT treatment: regression is instantly, and almost complete at 18 days. Relapse is apparent 22 days after 4-OHT treatment (see also Fig. S5a). **c)** Average tumor volume plotted against time. Data are averages of different tumors (WT/WT (n=8), TA/TA (n=10), TA/KD (n=7), KD/KD (n=19). Volumes are relative to normalized volume at the start of 4-OHT treatment. Error bars indicate ±SD (see also Fig. S5b). **d)** Mutant expression in tumors from *CiMKi^TA/TA^, CiMKi^TA/KD^,* and *CiMKi^KD/KD^* mice.

## Discussion

We recently showed that the effects of CIN on intestinal tumorigenesis strongly depend on its degree and location along the intestinal tract (34). Here we assessed the effects of various CIN levels on tumor growth and regression dynamics, using the classic two-stage DMBA/TPA skin carcinogenesis model that enables facile longitudinal monitoring. In line with our study of intestinal tumorigenesis (34), the moderate level of CIN had the most profound impact, causing accelerated onset, increased frequency and enhanced growth rates of the tumors. High and very high CIN accelerated tumor onset and growth but did not enhance the number of tumors, while low and very low CIN only enhanced the latter. Our present work again emphasizes the relevance of tissue and context on the impact of CIN on tumorigenesis: high CIN (TA/KD) does not promote tumorigenesis in the small intestine of *Apc^Min/+^* mice but does so in the large intestine (34), and, as we show here, CIN is pro-tumorigenic also in the skin after DMBA and DMBA/TPA application. Moreover, while moderate and higher CIN levels alone (no other oncogenic treatment or mutation) could promote intestinal tumorigenesis, none of the CIN levels could do so in the skin within the observed duration of 45 weeks (not shown).

What causes these different effects of CIN on tumorigenesis? Our data point to cell-intrinsic and -extrinsic effects that are differentially induced by the various degrees of CIN. Besides copy number gains of chromosomes 6 and 7, common to the no-CIN tumors, tumors of all CIN genotypes were found to have other gains and losses, indicating that CIN increases the chance for the epithelial cells to reach an oncogenic karyotype, leading to a higher probability of tumor initiation and thus higher numbers of tumors. On the other hand, high and very high CIN tissue showed signs of increased cell death, immune evasion, and regeneration shortly after induction, suggesting additional cell-extrinsic effects of these high CIN levels (fig 8). This was further supported by the surprising finding that the fast-growing tumors after moderate/high CIN induction consisted of cells with low or very low CIN genotypes. This was true for tumors of the skin, the breast, the large intestine and the mesentery, suggesting a more universal principle of non-cell autonomous tumor promotion by high CIN. The fact that the tumors were uniformly of the same (very) low CIN genotype instead of mixed alleles shows that they most probably arose from a single clone. This is in line with recent data showing monoclonal origins of DMBA/TPA-induced SCC in mice (52). Given the observed high efficiency of recombination in vivo, assessed soon after induction, these data suggest that the fast disappearance of high CIN cells greatly stimulated survival and proliferation of very rare low CIN cells. The speed at which those rare cells grew to become a tumor is a striking example of strong non-cell autonomous effects of high CIN on tumor formation. While we have not fully uncovered the mechanisms, our data suggest that inflammation does not play a major role: immune cells were mostly excluded from the tissue following CIN induction, and the NSAID carprofen did not substantially affect the impact of CIN on tumor promotion. This was unexpected, as CIN has known pro-inflammatory effects and we show it can supplant TPA as the promoting agent following initiation by DMBA. CIN and TPA therefore likely have other overlapping tumor-promoting mechanisms. The exact mechanism of the strong non-cell-autonomous oncogenic property of high CIN will be of great interest to uncover, as it might guide new therapeutic strategies.

**Figure 8.**
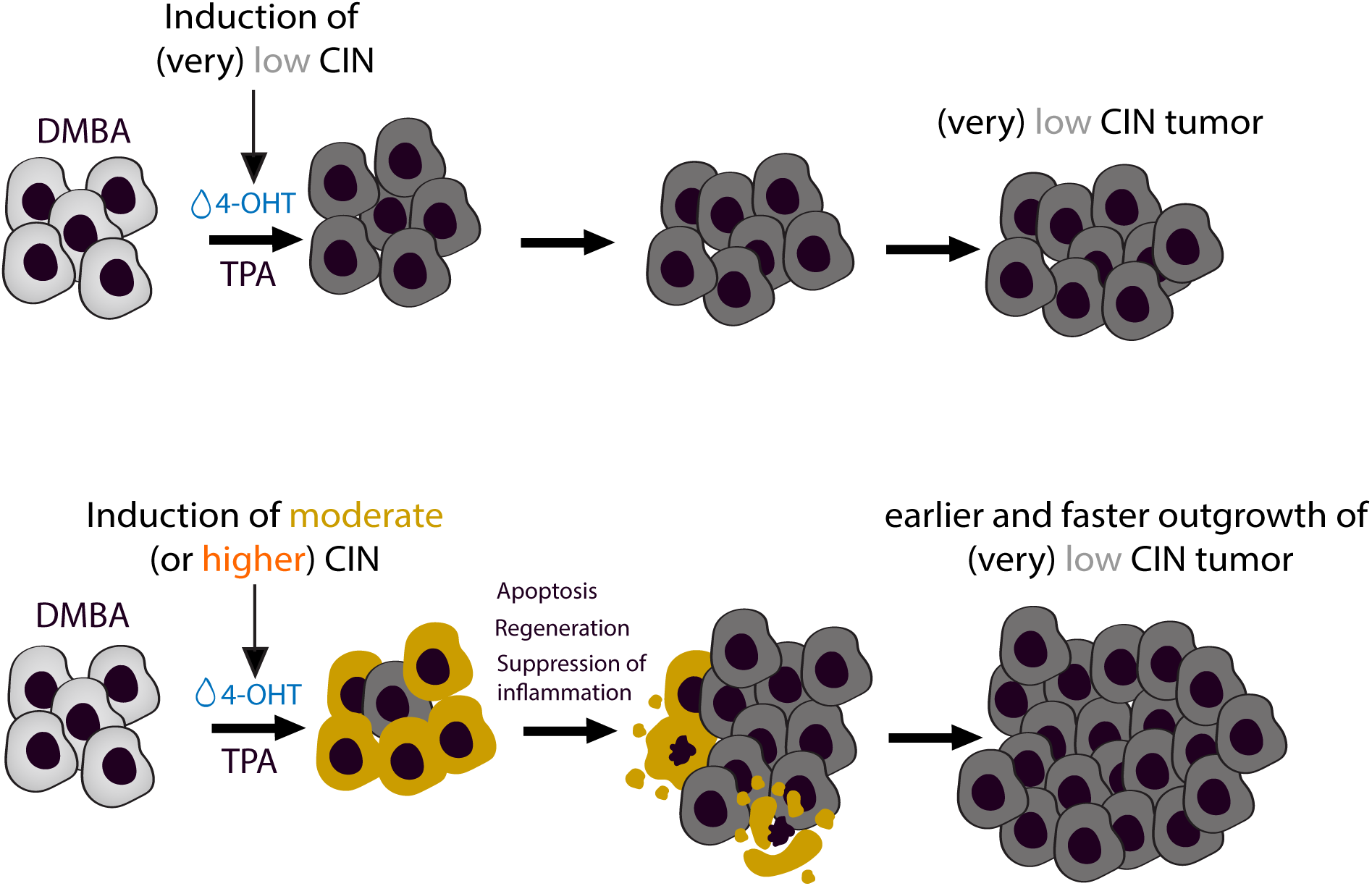
Model for cell-extrinsic effects of high CIN levels in tumorigenesis. Moderate or higher CIN levels rapidly disappear from the tissue, possibly due to unviable levels of aneuploidy, but while doing so, they create a microenvironment ideal for rare low CIN cells to thrive. They induce a regenerative response accompanied by lack of immune cells and inflammation, which possibly explains the accelerated onset and fast growth but lower number of tumors by (very) high CIN. Low and very low CIN lack the cell-extrinsic effect, resulting only in increased tumor frequency.

Taken together, our data suggest that the differential effects of CIN on tumor initiation and growth originate from both cell-intrinsic and cell-extrinsic effects: the oncogenic cell-intrinsic effects of CIN (e.g. accelerated evolution of oncogenic karyotypes) are most efficiently sustained by low or very low CIN, causing increased initiation frequency; while oncogenic cell-extrinsic effects of CIN (e.g. a hyper-regeneration response) are most efficiently created by higher CIN, speeding up growth. These effects are combined most optimally in the moderate CIN cells, causing acceleration of tumor onset, increase in initiation and increase in growth. Low and very low CIN lack the cell-extrinsic effect, resulting in predominantly accelerated onset. High and very high CIN cells eventually succumb, possibly due to unviable levels of aneuploidy, but leave a microenvironment ideal for rare low CIN cells to proliferate. This explains the accelerated onset and fast growth but lower number of tumors by (very) high CIN. It will be of great interest to untangle how the higher CIN levels cause the regenerative response, whether it is acute or chronic, and which features of the response have which effects on tumor initiation and growth.

The skin tumorigenesis model allowed for the monitoring of tumor size before and after CIN induction, which mimics therapeutic application of CIN-enhancing drugs such as microtubule poisons or inhibitors of mitotic kinases or kinesins (e,g, Aurora, MPS1, PLK1, Kif18A or Kif11). We observed fast regression especially upon high and very high CIN induction. However, tumors relapsed quickly. Strikingly, the high CIN appeared to have promoted fast growth of rare low CIN cells, similar to the effects of high CIN during tumor initiation. Caution is therefore warranted when designing CIN-based therapeutic strategies. Optimizing the potential success of such strategies will require additional research into the mechanisms of the non-cell-autonomous effects of high CIN on tumor relapse.

## Methods

### Mice: strains, experiments, and analysis

Animal experiments were approved by Animal Experimental Committee and the Dutch Central Authority for Scientific Procedures on Animals (CCD) and carried out in accordance with DIRECTIVE 2010/63/EU OF THE EUROPEAN PARLIAMENT AND OF THE COUNCIL. Animals were bred and housed under standard conditions (humidity 50-60%, 22-23°C, inverted 12/12 hour light/dark cycle, water and standard chow ad libitum) at the animal facility of the Gemeenschappelijk Dieren Laboratorium (GDL), Utrecht, the Netherlands, and the Hubrecht animal facility. Genetically modified mouse strains used in this study include: Cre-inducible-Mps1-Knock-in-TA (CiMKi^TA^) and Cre-inducible-Mps1-Knock-in-KD (CiMKi^KD^) (34) and available at JAX® (mice (B6.129P2(SJL)-*Ttk^tm1.1Kops^*/JlmaJ (stock #035136) and B6.129P2(SJL)-*Ttk^tm2.1Kops^*/JlmaJ (stock#035137), respectively), Rosa26-CreER^T2^ (purchased from JAX® mice (B6.129Gt(ROSA)26Sor^tm1(cre/ERT2)Tyj^/J, stock#008463)), and p53^LoxP^ (purchased from JAX® mice (B6.129P2-*Trp53^tm1Brn^*/J, stock#008462)). All mice were maintained in C57BL/6 background. For experiments involving p53 knock-out mice, *CiMKi; Rosa26-CreERT2; p53^+/+^* and *CiMKi; Rosa26-CreERT2; p53^+/Flox^* mice were generated. CIN and floxing of the p53 allele were induced by intraperitoneal injection with Tamoxifen (1 mg dissolved in corn oil; Sigma, C8267) when mice were five weeks old. Mice were checked for general health and discomfort (according to the guidelines of the Dutch Code of Practice Animals in Cancer Research), weighed and palpated for the presence of tumors once a week. Mice were killed when a clear tumor could be palpated or when a humane endpoint was reached, or after maximally 104 weeks. Whole body dissection was performed, and tissues and tumors were divided in two; and formalin fixed and embedded in paraffin, and snap-frozen in liquid nitrogen for further analyses. All mice were genotyped using standard PCR and targeted sequencing procedures (JAX® Mice and (34), also see Supplementary Table 1).

### Skin Tumorigenesis

For skin tumorigenesis experiments *CiMKi;Rosa26-CreERT2* of all CiMKi combinations were generated. To induce tumors, the classical DMBA/TPA skin carcinogenesis protocol (40) was followed, in combination with induction of various levels of CIN: back skin of 5- to 6-weeks old female mice was shaven and topically treated with a single dose of 7,12-dimethylbenz[a]-anthracene (DMBA) (800 nmol/0.2 ml acetone, Sigma). One week later, biweekly topical treatment of the back skin with 12-O-tetradecanoylphorbol-13-acetate (TPA) (4 µg/0.2 ml acetone, Sigma) started, for the duration of the experiment. CIN was induced ten days after DMBA treatment by topical treatment on the back skin with 4-hydroxytamoxifen (4-OHT) (1 mg/0.2 ml DMSO, Sigma H6278) (five singles doses on consecutive days). To induce CIN at a later timepoint in established tumors (Figures 7, S5), 4-OHT was applied in a similar way when tumors were 100-200mm^3^ in size. In case mice were treated with Carprofen (0,06 mg/ml, ProduLab Pharma BV) was added to the drinking water for the duration of the experiment. To monitor tumor growth, the backs of the mice were shaven as often as necessary and tumor volume was calculated by taking length and width measurements with a caliper every week. The average of these values was divided by two and taken as radius (volume = 4/3*π*\*r^3^). Mice were euthanized when total tumor volume exceeded 1500 mm^3^, or maximally when 45 weeks in experiment. At dissection, tumors, normal skin and internal organs were isolated and divided to be snap-frozen or fixed in formalin. For identification and assessment of tumors 4-μm sections of paraffin-embedded tissue were cut and stained with hematoxylin/eosin (H&E). Grading of tumors was done by pathologists from the Dutch Molecular Pathology Centre.

### Establishment and culture of skin organoids

3D organoids cultures were derived from back skins of six-to-twelve weeks-old, culled *CiMKi;Apc^Min/+^;Rosa26-CreER^T2^*mice as described in (39). Organoids were cultured in advanced DMEM/F12 (Gibco) supplemented with penicillin/ streptomycin (100 U/L; Gibco), Hepes, (10 mM; Gibco), GlutaMAX (1×; Gibco), B27 supplement (50× stock; Gibco), N-Acetylcysteine-1 (1 mM; SigmaAldrich), Noggin-conditioned medium (5%, kind gift from the Clevers group), R-spondin 1-conditioned medium (5%, kind gift from the Clevers group), acidic FGF1 (100 μg/mL; Peprotech), Heparin (0.2%; Stemcell Technologies), Forskolin (10 ng/mL; Tocris), Rho kinase inhibitor (Y-27632; 10 μM; Sigma-Aldrich), and Primocin (500× stock; Invivogen).For passaging, organoids were sheared by repetitive pipetting and re-plated in Matrigel in a pre-warmed 24-well plate.

### Assessment of CiMKi mutant allele expression

To confirm recombination of the CiMKi alleles and mutant expression, RNA was isolated from tumors, normal skin or skin organoids using the quick RNA kit (Zymo Research). cDNA was prepared using standard procedures, subjected to PCR and subsequently sequenced to determine the presence of T649A or D637A. For primers see Supplementary Table 1.

### Histology and Immunohistochemistry

4-5 μm sections of paraffin-embedded tissue were cut and stained with hematoxylin/eosin (H&E), or Masson’s according to standard protocols. Primary antibodies used for immunohistochemistry were against ki67 (ThermoFisher, RM-9106, ARS pH9, 1:50 and 3-hour incubation), PCNA (Clone PC10; Millipore MAB424,No AR, 1:100, overnight incubation), SMA (Sigma, A5228 clone 1A4, No AR, 1:4000, 1hr incubation), Gli1 (Thermofisher, MA5-32553, AR EDTA ph9, 1:150, overnight incubation), Gli2 (Thermofisher, CF804549, AR EDTA ph9, 1:150, overnight incubation), CD45 (BioLegend, 103102, AR Citrate pH6, 1:500, overnight incubation), and Cleaved Caspase-3 (Cell signaling, Cleaved Caspase 3 (D175); 9664S,AR EDTA pH9 ,1:1000, 1hr incubation). Secondary antibodies were anti-Rabbit Envision-HRP (DAKO, one hour), and anti-Mouse Envision-HRP (DAKO, one hour). Counterstaining was done with hematoxylin.

### Timelapse microscopy

To analyze chromosome segregation using fluorescence microscopy time-lapse imaging, single cells that were cryopreserved immediately after collection were thawed and transduced with an H2B-Neon expressing lentivirus (pLV-H2B-mNeon-puro (34)). For induction of CiMKi alleles organoids were treated with 1 μM 4-OHT for 56 hours. Organoids were seeded 24 hours before imaging in 8-chamber IBIDI slides and imaged with a confocal spinning disk (Nikon/Andor CSU-W1 with Borealis illumination), equipped with atmospheric and temperature control. Organoids were imaged in XYZT-mode (12 to 20 z-sections at 2.5 μm intervals, for 8 to 12 hours) at 37°C at 3-minute intervals, using a 30X silicon objective and an additional 1.5X lens in front of the CCD-camera. 3% 448nm laser and 50nm disk pinhole were used. Raw data were converted to videos using a customised ImageJ/Fiji macro(36, 53). Fidelity of all observed chromosome segregations was scored and categorized manually, in a blinded manner.

### Next Generation Sequencing

For bulk whole genome sequencing, genomic DNA was isolated from tumours and normal skin with the DNeasy Blood and Tissue kit (Qiagen) according to the manufacturer’s instruction. Sequencing and copy number analysis were done at the Utrecht Sequencing Facility (useq.nl).

### Bulk RNA sequencing

Back skin of *CiMKi;Rosa26-CreER^T2^* (no CIN, moderate (TA/TA) and high (TA/KD)) were treated with 4-hydroxytamoxifen (4-OHT) (1 mg/0.2 ml DMSO, Sigma H6278) (five singles doses on consecutive days) and were harvested 7 and 28 days after first application. RNA was isolated from the back skin of the mice using the Quick-RNA Tissue/Insect microprep kit (Zymo Research). A total of 500 ng RNA per sample was used as input material for the RNA sample preparations. RNA integrity number (RIN) was assessed using the RNA Nano 6000 Assay Kit of the Bioanalyzer 2100 system (Agilent Technologies, Santa Clara, CA) with a threshold of 7. Two samples were excluded (high CIN) because of poor RNA quality, therefore not all conditions have three biological replicates. Briefly, mRNA was purified from total RNA using NEXTFLEX Poly(A) Beads (Perkin elmer, NOVA-512993). Library preparation was executed using the kit (NEXTFLEX® Rapid Directional RNA Kit 2.0, NOVA-5198-03). Library quality was assessed on the Agilent Bioanalyzer 2100 system. The samples were barcoded using NEXTFLEX® Unique Dual Index Barcodes (Perkin elmer, NOVA-512920). The libraries were sequenced on Nextseq2000p2, single and paired-end reads were generated, at the USEQ (Utrecht Sequencing Facility, Useq.nl). Raw data (raw reads) of fastq format were processed at the USEQ in order to correct the raw counts for sequencing differences between samples. Intergroup differences were evaluated using the principal component analysis, which revealed that the primary source of variability in the data was the effect of CIN over time (Fig. S4a). After 7 days from CIN induction, high (TA/KD) CIN skin, but not moderate (TA/TA) CIN, clusters distantly from no CIN samples. However, after 28 days from CIN induction, both high (TA/KD) and moderate (TA/TA) CIN samples cluster distantly from no CIN condition. To find CIN-specific induced transcriptomic profile, we conducted pairwise comparisons between CIN and no CIN samples at both timepoints. differentially expressed genes (DEGs). Differential expression analysis was performed using DESeq2 R package (version 1.30.1). The significance threshold applied for DEGs was adjusted *P*-value/false discovery rate < 0.05 and |Log fold change| > 1. Gene ontology analysis was performed using the Metascape online tool (version 3.5). (54)

### Statistics and data reproducibility

For mouse tumorigenesis studies, a priori power analyses dictated the number of animals that were enrolled in each group to detect differences with 80% power and 95% confidence. Animals were randomized by combining various genotypes per cage. Treatments, tumor measurements, and analyses were done blinded. For mouse tumorigenesis and organoid studies, statistical analyses were done using GraphPad Prism. Data are presented as averages ± SD unless otherwise stated in legends. Statistical tests are indicated in the figure legends. All images and micrographs shown are representative for experimental groups of at least three individual mice in each experiment.

## Supporting information

Supplemental images and legends

## DATA AVAILABILITY

The WGS and RNAseq data have been deposited in the European Nucleotide Archive database under the accession code PRJEB100986. All the other data obtained for this study are available within the article and its supplementary information files and from the corresponding author upon request.

## Acknowledgments.

We thank the Hubrecht animal facility, the Hubrecht FACS facility, the Hubrecht imaging facility, Single Cell Discoveries, and the Utrecht Sequencing Facility (USEQ) for assistance with the experiments. We thank Kim Boonekamp and Kai Kretschmar for support with skin organoid cultures, Harry Begthel, Jeroen Korving and Jaimy Tatuhey for support with histology and stainings, and Dani Bodor for help with image processing. The Kops lab is part of Oncode Institute, which is partly funded by the Dutch Cancer Society (KWF Kankerbestrijding). This work was supported by the Dutch Cancer Society (grant numbers HUBR-2012-5321, HUBR-2012-5513, and KWF 10126) and by the European Research Council (ERC-SyG 855158).

## Author contributions

A.J., N.J. and G.J.P.L.K. designed the research. N.J. and S.L. analyzed and assembled the data. B.G., S.L. A.I.Q., A.J., W.H.M.H., B.E., N.C. and N.J. performed animal maintenance and experiments. B.G. and S.L. performed and analyzed scKaryo-seq and RNAseq. P.T. performed organoid experiments. S.Y., L.A.A.B. and A.d.B. performed pathology analyses. S.L., N.J. and G.J.P.L.K wrote the paper.

## Competing interests

The authors declare no competing interests.

## Notes

### Competing Interest Statement

The authors have declared no competing interest.

## References

1. van Jaarsveld RH, Kops G. Difference Makers: Chromosomal Instability versus Aneuploidy in Cancer. Trends in cancer. 2016;2(10):561–71.

2. Heim SaM, F. Cancer Cytogenetics: Chromosomal and Molecular Genetic Aberrations of Tumor Cells. 1995;2nd Edition, Wiley-Liss, Inc., New York.

3. Kristin A. Knouse TD, , Stephen J. Elledge,, and Angelika Amon. Aneuploidy in Cancer: Seq-ing Answers to Old Questions. Annual Review of Cancer Biology 2017;Vol. 1:335–354.

4. Bolhaqueiro ACF, Ponsioen B, Bakker B, Klaasen SJ, Kucukkose E, van Jaarsveld RH, et al. Ongoing chromosomal instability and karyotype evolution in human colorectal cancer organoids. Nature genetics. 2019;51(5):824–34.

5. Bach DH, Zhang W, Sood AK. Chromosomal Instability in Tumor Initiation and Development. Cancer research. 2019;79(16):3995–4002.

6. Serçin Ö, Larsimont JC, Karambelas AE, Marthiens V, Moers V, Boeckx B, et al. Transient PLK4 overexpression accelerates tumorigenesis in p53-deficient epidermis. Nature cell biology. 2016;18(1):100–10.

7. Foijer F, Xie SZ, Simon JE, Bakker PL, Conte N, Davis SH, et al. Chromosome instability induced by Mps1 and p53 mutation generates aggressive lymphomas exhibiting aneuploidy-induced stress. Proceedings of the National Academy of Sciences of the United States of America. 2014;111(37):13427–32.

8. Baker DJ, Jin F, Jeganathan KB, van Deursen JM. Whole chromosome instability caused by Bub1 insufficiency drives tumorigenesis through tumor suppressor gene loss of heterozygosity. Cancer cell. 2009;16(6):475–86.

9. Michel LS, Liberal V, Chatterjee A, Kirchwegger R, Pasche B, Gerald W, et al. MAD2 haplo-insufficiency causes premature anaphase and chromosome instability in mammalian cells. Nature. 2001;409(6818):355–9.

10. Weaver BA, Silk AD, Montagna C, Verdier-Pinard P, Cleveland DW. Aneuploidy acts both oncogenically and as a tumor suppressor. Cancer cell. 2007;11(1):25–36.

11. Bakhoum SF, Ngo B, Laughney AM, Cavallo JA, Murphy CJ, Ly P, et al. Chromosomal instability drives metastasis through a cytosolic DNA response. Nature. 2018;553(7689):467–72.

12. Levine MS, Bakker B, Boeckx B, Moyett J, Lu J, Vitre B, et al. Centrosome Amplification Is Sufficient to Promote Spontaneous Tumorigenesis in Mammals. Developmental cell. 2017;40(3):313–22.e5.

13. Sotillo R, Hernando E, Díaz-Rodríguez E, Teruya-Feldstein J, Cordón-Cardo C, Lowe SW, Benezra R. Mad2 overexpression promotes aneuploidy and tumorigenesis in mice. Cancer cell. 2007;11(1):9–23.

14. Rao CV, Yang YM, Swamy MV, Liu T, Fang Y, Mahmood R, et al. Colonic tumorigenesis in BubR1+/-ApcMin/+ compound mutant mice is linked to premature separation of sister chromatids and enhanced genomic instability. Proceedings of the National Academy of Sciences of the United States of America. 2005;102(12):4365–70.

15. Ricke RM, Jeganathan KB, van Deursen JM. Bub1 overexpression induces aneuploidy and tumor formation through Aurora B kinase hyperactivation. The Journal of cell biology. 2011;193(6):1049–64.

16. Trakala M, Aggarwal M, Sniffen C, Zasadil L, Carroll A, Ma D, et al. Clonal selection of stable aneuploidies in progenitor cells drives high-prevalence tumorigenesis. Genes & development. 2021;35(15-16):1079–92.

17. Shoshani O, Bakker B, de Haan L, Tijhuis AE, Wang Y, Kim DH, et al. Transient genomic instability drives tumorigenesis through accelerated clonal evolution. Genes & development. 2021;35(15-16):1093–108.

18. Iwanaga Y, Chi YH, Miyazato A, Sheleg S, Haller K, Peloponese JM, Jr., et al. Heterozygous deletion of mitotic arrest-deficient protein 1 (MAD1) increases the incidence of tumors in mice. Cancer research. 2007;67(1):160–6.

19. Girish V, Lakhani AA, Thompson SL, Scaduto CM, Brown LM, Hagenson RA, et al. Oncogene-like addiction to aneuploidy in human cancers. Science (New York, NY). 2023;381(6660):eadg4521.

20. Zhang CZ, Spektor A, Cornils H, Francis JM, Jackson EK, Liu S, et al. Chromothripsis from DNA damage in micronuclei. Nature. 2015;522(7555):179–84.

21. Ly P, Teitz LS, Kim DH, Shoshani O, Skaletsky H, Fachinetti D, et al. Selective Y centromere inactivation triggers chromosome shattering in micronuclei and repair by non-homologous end joining. Nature cell biology. 2017;19(1):68–75.

22. Ly P, Brunner SF, Shoshani O, Kim DH, Lan W, Pyntikova T, et al. Chromosome segregation errors generate a diverse spectrum of simple and complex genomic rearrangements. Nature genetics. 2019;51(4):705–15.

23. Papathanasiou S, Mynhier NA, Liu S, Brunette G, Stokasimov E, Jacob E, et al. Heritable transcriptional defects from aberrations of nuclear architecture. Nature. 2023;619(7968):184–92.

24. Shoshani O, Brunner SF, Yaeger R, Ly P, Nechemia-Arbely Y, Kim DH, et al. Chromothripsis drives the evolution of gene amplification in cancer. Nature. 2021;591(7848):137–41.

25. Li J, Hubisz MJ, Earlie EM, Duran MA, Hong C, Varela AA, et al. Non-cell-autonomous cancer progression from chromosomal instability. Nature. 2023;620(7976):1080–8.

26. Bakhoum SF, Cantley LC. The Multifaceted Role of Chromosomal Instability in Cancer and Its Microenvironment. Cell. 2018;174(6):1347–60.

27. Hong C, Schubert M, Tijhuis AE, Requesens M, Roorda M, van den Brink A, et al. cGAS-STING drives the IL-6-dependent survival of chromosomally instable cancers. Nature. 2022;607(7918):366–73.

28. van den Brink A, Suárez Peredo Rodríguez MF, Foijer F. Chromosomal instability and inflammation: a catch-22 for cancer cells. Chromosome research : an international journal on the molecular, supramolecular and evolutionary aspects of chromosome biology. 2023;31(3):19.

29. Wang RW, Viganò S, Ben-David U, Amon A, Santaguida S. Aneuploid senescent cells activate NF-κB to promote their immune clearance by NK cells. EMBO reports. 2021;22(8):e52032.

30. Davoli T, Uno H, Wooten EC, Elledge SJ. Tumor aneuploidy correlates with markers of immune evasion and with reduced response to immunotherapy. Science (New York, NY). 2017;355(6322).

31. Burrell RA, McGranahan N, Bartek J, Swanton C. The causes and consequences of genetic heterogeneity in cancer evolution. Nature. 2013;501(7467):338–45.

32. Sansregret L, Vanhaesebroeck B, Swanton C. Determinants and clinical implications of chromosomal instability in cancer. Nature reviews Clinical oncology. 2018;15(3):139–50.

33. Lukow DA, Sheltzer JM. Chromosomal instability and aneuploidy as causes of cancer drug resistance. Trends in cancer. 2022;8(1):43–53.

34. Hoevenaar WHM, Janssen A, Quirindongo AI, Ma H, Klaasen SJ, Teixeira A, et al. Degree and site of chromosomal instability define its oncogenic potential. Nature communications. 2020;11(1):1501.

35. Silk AD, Zasadil LM, Holland AJ, Vitre B, Cleveland DW, Weaver BA. Chromosome missegregation rate predicts whether aneuploidy will promote or suppress tumors. Proceedings of the National Academy of Sciences of the United States of America. 2013;110(44):E4134–41.

36. Drost J, van Jaarsveld RH, Ponsioen B, Zimberlin C, van Boxtel R, Buijs A, et al. Sequential cancer mutations in cultured human intestinal stem cells. Nature. 2015;521(7550):43–7.

37. Thompson LL, Jeusset LM, Lepage CC, McManus KJ. Evolving Therapeutic Strategies to Exploit Chromosome Instability in Cancer. Cancers. 2017;9(11).

38. Bakhoum SF, Compton DA. Chromosomal instability and cancer: a complex relationship with therapeutic potential. The Journal of clinical investigation. 2012;122(4):1138–43.

39. Boonekamp KE, Kretzschmar K, Wiener DJ, Asra P, Derakhshan S, Puschhof J, et al. Long-term expansion and differentiation of adult murine epidermal stem cells in 3D organoid cultures. Proceedings of the National Academy of Sciences of the United States of America. 2019;116(29):14630–8.

40. Abel EL, Angel JM, Kiguchi K, DiGiovanni J. Multi-stage chemical carcinogenesis in mouse skin: fundamentals and applications. Nature protocols. 2009;4(9):1350–62.

41. Fujiki H, Suganuma M, Yoshizawa S, Kanazawa H, Sugimura T, Manam S, et al. Codon 61 mutations in the c-Harvey-ras gene in mouse skin tumors induced by 7,12-dimethylbenz[a]anthracene plus okadaic acid class tumor promoters. Molecular carcinogenesis. 1989;2(4):184–7.

42. Rachana Garg AGR, Girish B. Maru. Curcumin decreases 12-O - tetradecanoylphorbol-13-acetate-induced protein kinase C translocation to modulate downstream targets in mouse skin. Carcinogenesis. 2008;29(6):Pages 1249–57.

43. C. Marcelo Aldaz DT, Fernando Larcher, Thomas J. Slaga, Claudio J. Conti. Sequential trisomization of chromosomes 6 and 7 in mouse skin premalignant lesions. Molecular carcinogenesis. 1989;2(1):Pages 22–6.

44. To MD, Quigley DA, Mao JH, Del Rosario R, Hsu J, Hodgson G, et al. Progressive genomic instability in the FVB/Kras(LA2) mouse model of lung cancer. Molecular cancer research : MCR. 2011;9(10):1339–45.

45. Rundhaug JE, Fischer SM. Molecular mechanisms of mouse skin tumor promotion. Cancers. 2010;2(2):436–82.

46. Ingrao JC, Johnson R, Tor E, Gu Y, Litman M, Turner PV. Aqueous stability and oral pharmacokinetics of meloxicam and carprofen in male C57BL/6 mice. Journal of the American Association for Laboratory Animal Science : JAALAS. 2013;52(5):553–9.

47. Coussens LM, Werb Z. Inflammation and cancer. Nature. 2002;420(6917):860–7.

48. Neagu M, Constantin C, Caruntu C, Dumitru C, Surcel M, Zurac S. Inflammation: A key process in skin tumorigenesis. Oncology letters. 2019;17(5):4068–84.

49. Maru GB, Gandhi K, Ramchandani A, Kumar G. The role of inflammation in skin cancer. Advances in experimental medicine and biology. 2014;816:437–69.

50. Love MI, Huber W, Anders S. Moderated estimation of fold change and dispersion for RNA-seq data with DESeq2. Genome Biology. 2014;15(12):550.

51. Hosea R, Hillary S, Naqvi S, Wu S, Kasim V. The two sides of chromosomal instability: drivers and brakes in cancer. Signal Transduction and Targeted Therapy. 2024;9(1):75.

52. Reeves MQ, Kandyba E, Harris S, Del Rosario R, Balmain A. Multicolour lineage tracing reveals clonal dynamics of squamous carcinoma evolution from initiation to metastasis. Nature cell biology. 2018;20(6):699–709.

53. Verissimo CS, Overmeer RM, Ponsioen B, Drost J, Mertens S, Verlaan-Klink I, et al. Targeting mutant RAS in patient-derived colorectal cancer organoids by combinatorial drug screening. eLife. 2016;5.

54. Zhou Y, Zhou B, Pache L, Chang M, Khodabakhshi AH, Tanaseichuk O, et al. Metascape provides a biologist-oriented resource for the analysis of systems-level datasets. Nature communications. 2019;10(1):1523.

