## Supplemental images and legends for "Non-cell-autonomous mechanisms of tumor initiation and relapse by chromosomal instability"

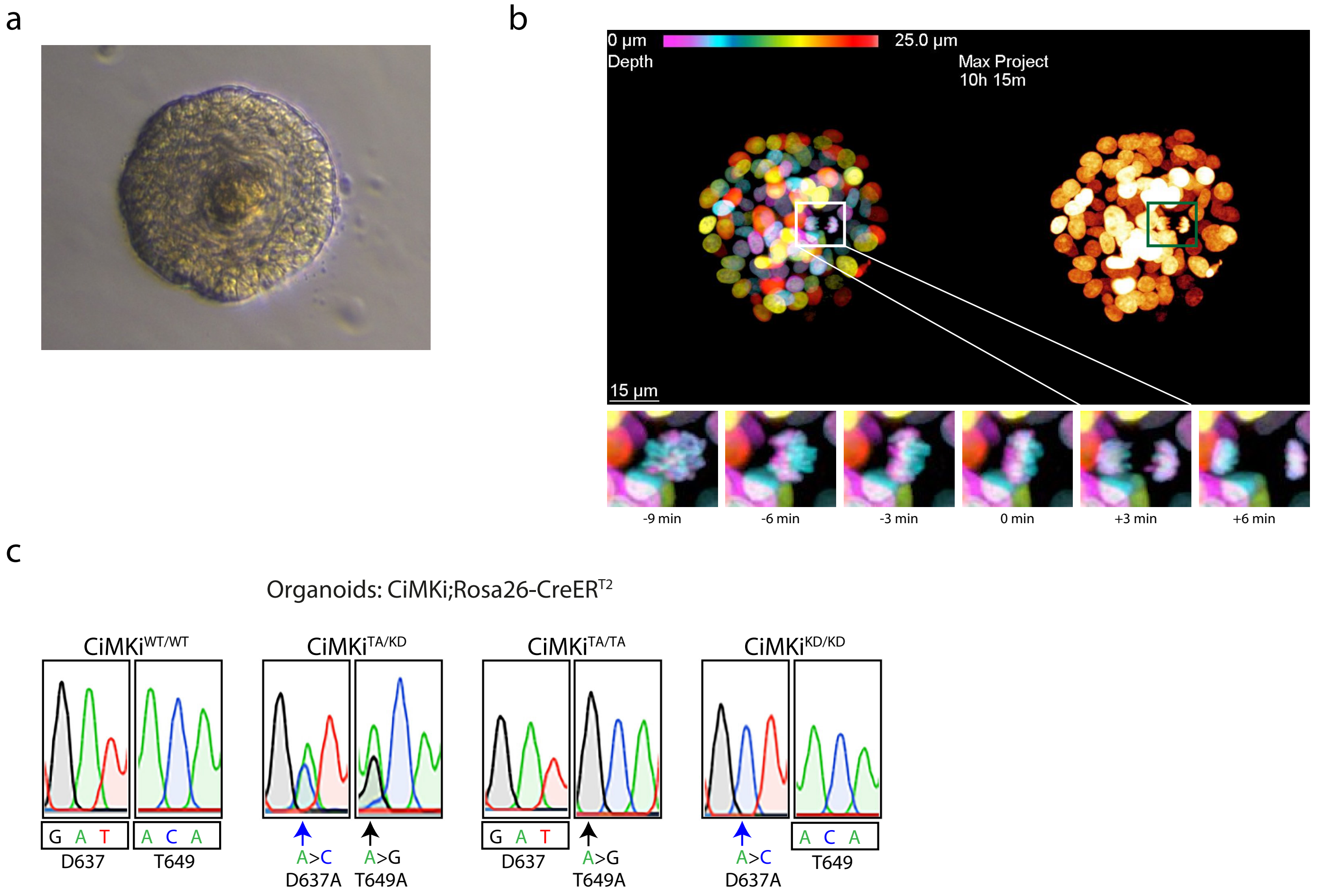
a
b
0 µm
25.0 µm
Depth
Max Project
10h 15m
15 µm
-9 min
-6 min
-3 min
0 min
+3 min
+6 min
c
Organoids: CiMKi;Rosa26-CreERT2
CiMKiWT/WT
G A T
D637
A C A
T649
CiMKiTA/KD
A>C
D637A
A>G
T649A
CiMKiTA/TA
G A T
D637
A>G
T649A
CiMKiKD/KD
A>C
D637A
A C A
T649

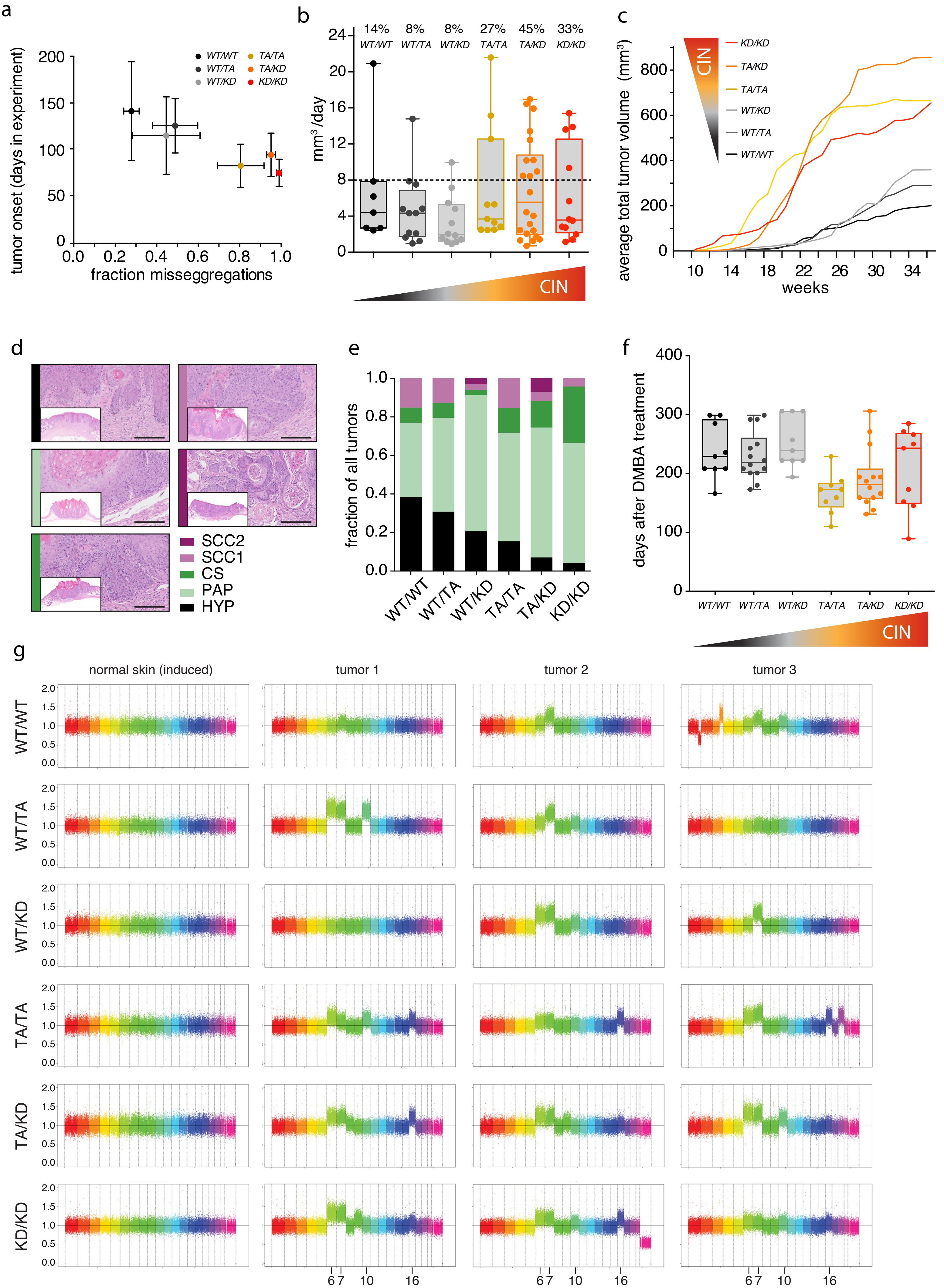


Lafirenze, van Gerwen et al, Figure S2

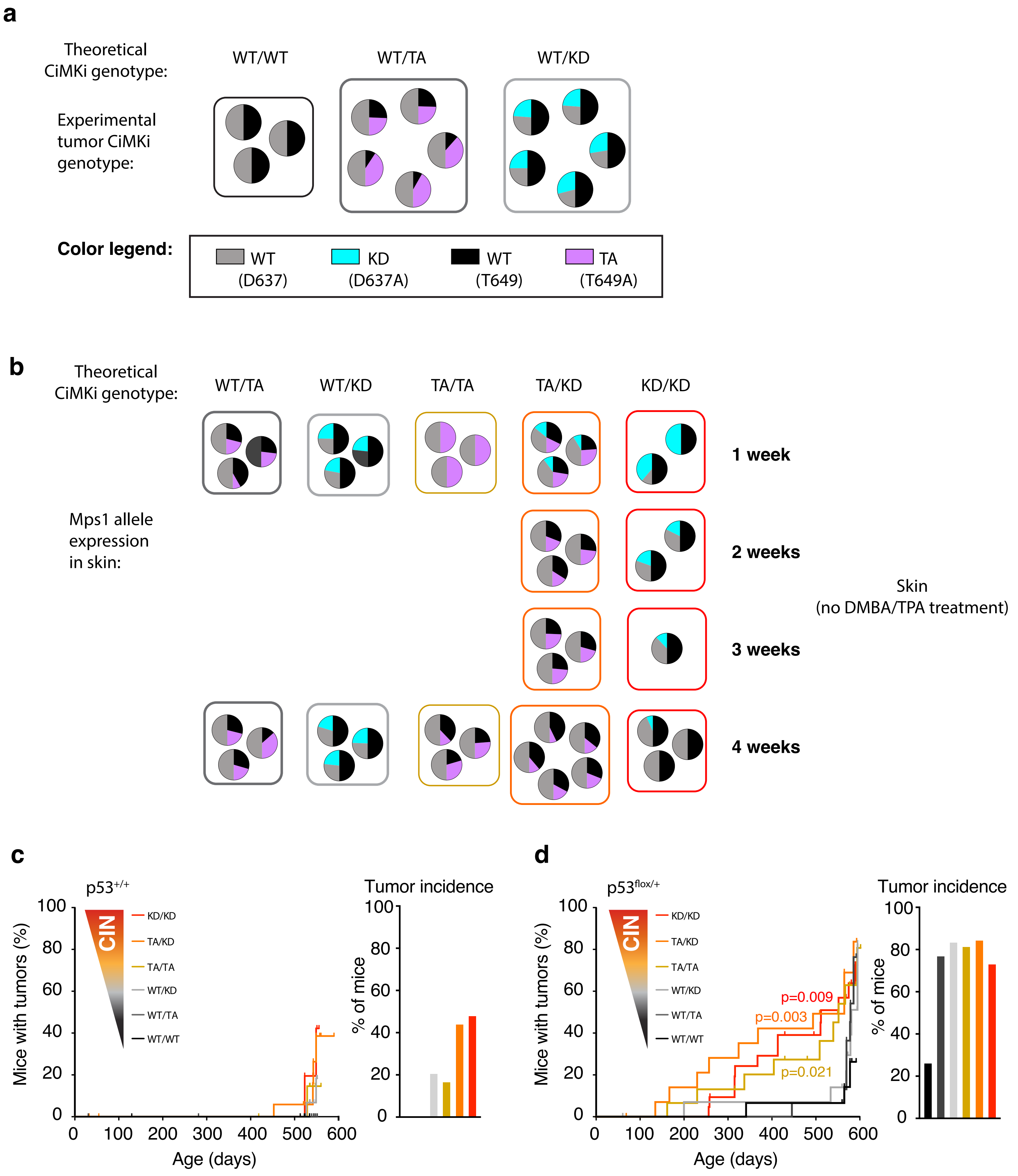


Lafirenze, van Gerwen et al, Figure S3

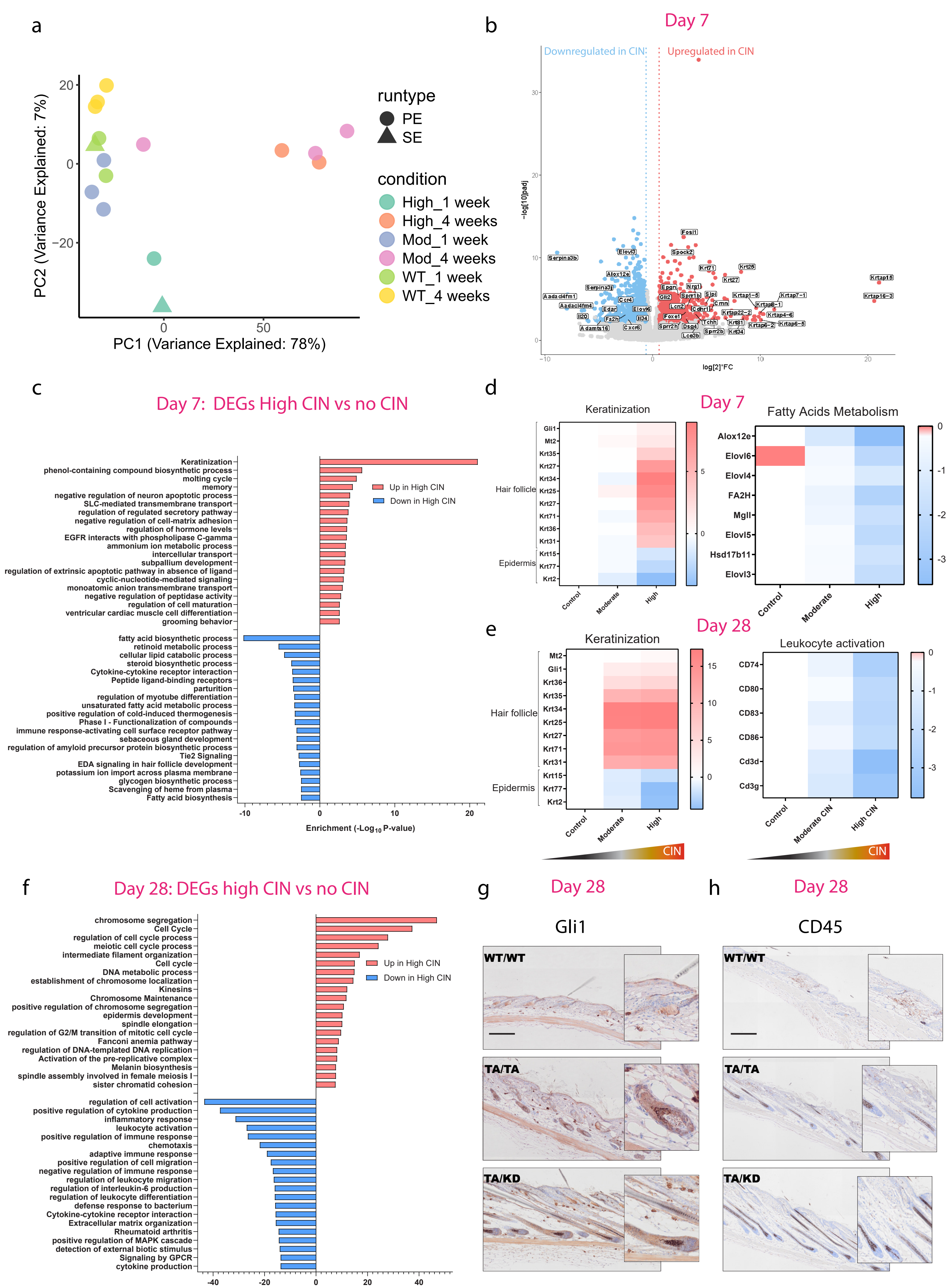


Lafirenze, van Gerwen et al, Figure S4

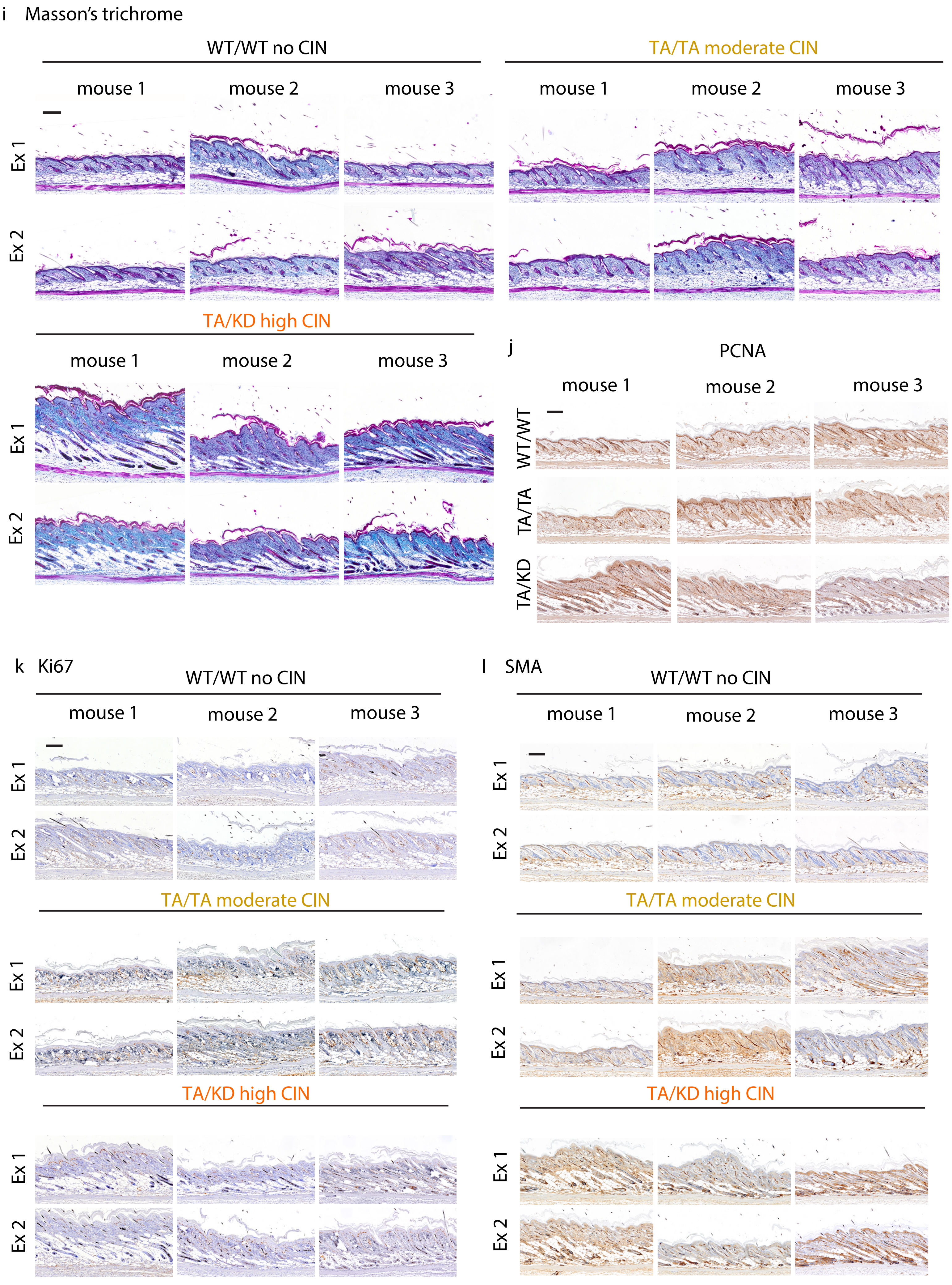


Lafirenze, van Gerwen et al., Figure S4i-l

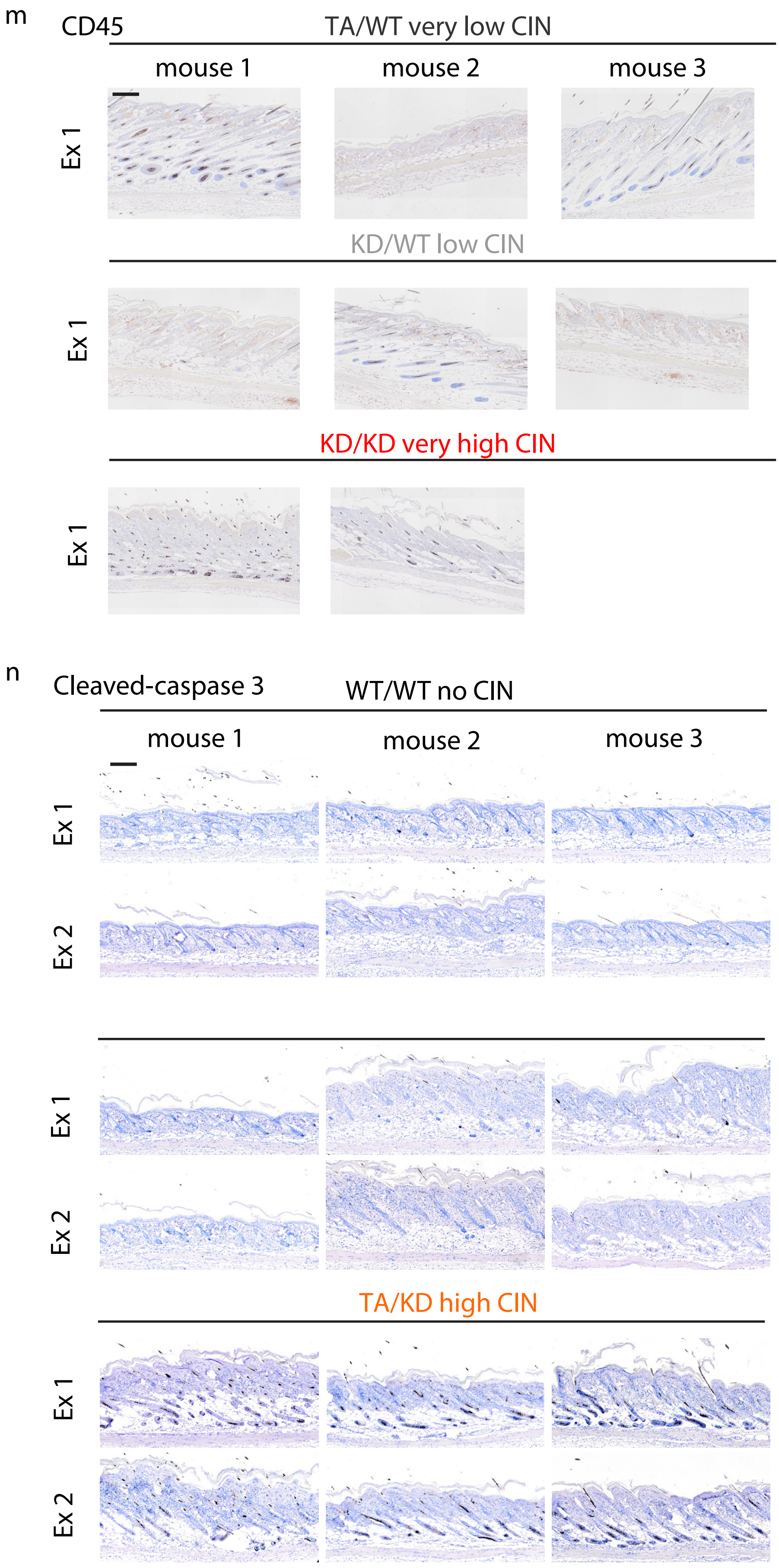


Lafirenze, van Gerwen et al., Figure S4m,n

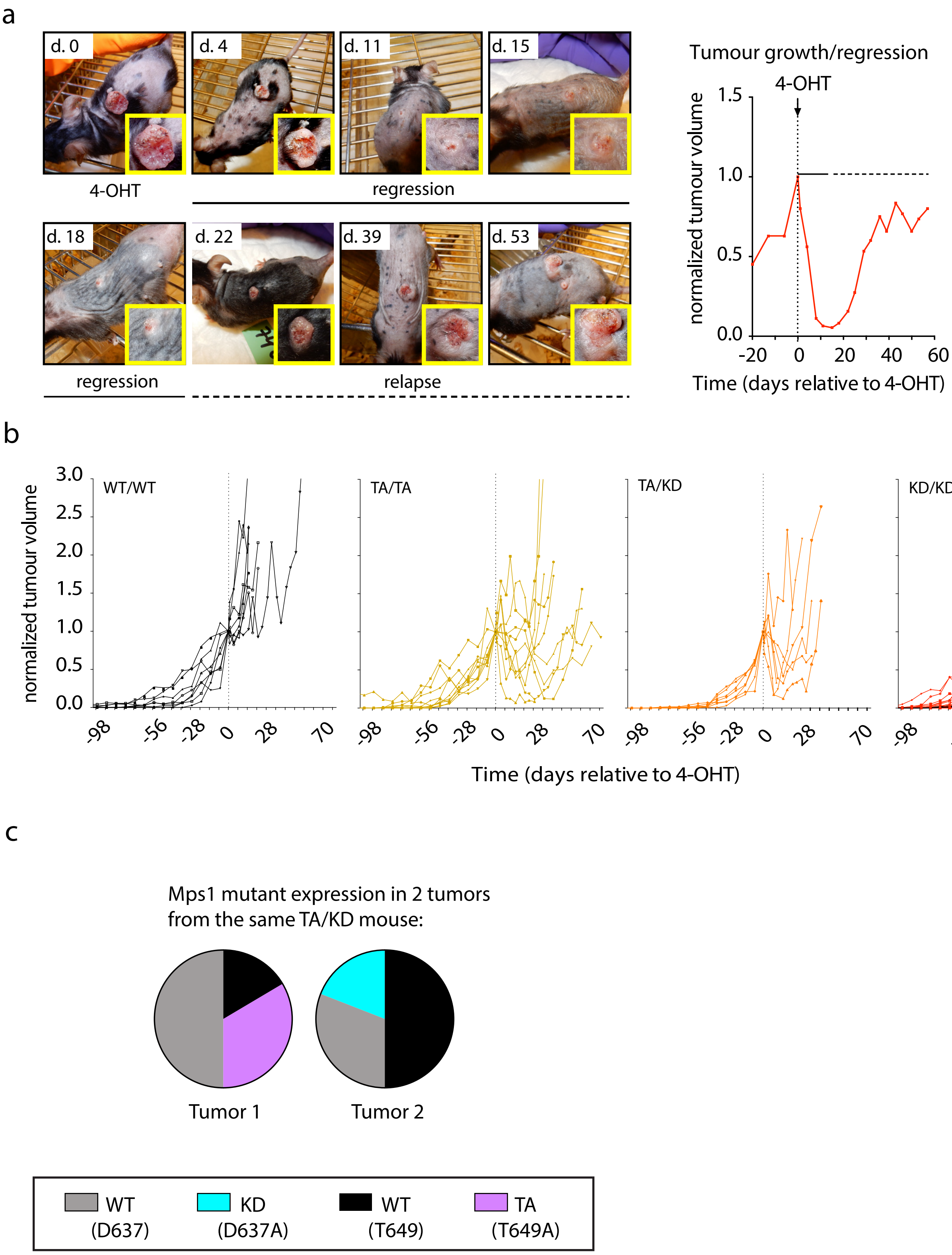


Lafirenze, van Gerwen et al., Figure S5

**Supplementary Figure legends and Supplementary table 1**

**S1.**

**a)** Image showing morphology (phase-contrast) of uninduced organoids derived from back skin of *CiMKi;Rosa26-CreER$^{T2}$* mice. **b)** Stills from timelapse movies of skin organoids of the *CiMKi;Rosa26-CreER$^{T2}$* genotypes 56 hours after 4-OHT addition. DNA is visualized by H2B-mNeon. Insets show 3-minute timepoints before and after metaphase (0 min.). **c)** Sequencing chromatograms of reverse transcribed RNA show efficient expression of both D637A (A to C) or T649A (A to G) CiMKi alleles in skin organoids of the *CiMKi;Rosa26-CreER$^{T2}$* genotypes 56 hours after 4-OHT addition.

**S2.**

**a)** Correlation plot of tumor onset in *CiMKi;Rosa26-CreER$^{T2}$* mice against the fraction of total missegregations in the *CiMKi* genotypes as measured in *CiMKi* organoids. Spearman correlation = -0886; p= 0.033). **b)** Box and whisker plot showing tumor growth rates for all tumors. Each data point represents one tumor. Boxes indicate 25$^{th}$ to 75$^{th}$ percentiles with median and error bars indicate min to max values. Above each box, the fraction of tumors that on average grew more than 8 mm$^3$ per day (=75$^{th}$ percentile of wild-type control) after exceeding 100 mm$^3$ is indicated. **c)** Average total tumor volume per mouse in time in *CiMKi;Rosa26-CreER$^{T2}$* mice. **d)** H&E examples of the various stages of skin tumorigenesis in DMBA/TPA promoted tumorigenesis in *CiMKi;Rosa26-CreER$^{T2}$* mice. Scale bars 100 µm, insets show zoom-out overviews of whole lesions. Classifications: hyperplastic tissue (HYP), papilloma (PAP), carcinoma in situ (CS), squamous cell carcinoma 1 and 2 (SCC1 and SCC2). **e)** The average number and fraction of the various stages of lesions per mouse in *CiMKi;Rosa26-CreER$^{T2}$* mice at the moment of sacrifice, classified as in (d). **f)** Time of sacrifice of *CiMKi;Rosa26-CreER$^{T2}$* mice. Boxes indicate 25$^{th}$ to 75$^{th}$ percentiles with median and error bars indicate min to max values. **g)** Rainbow plot showing deviations from the 2n state of individual chromosomes (whole genome bulk DNA sequencing), in normal skin and three individual tumors of all CiMKi genotypes.

**S3**

**a)** Mutant expression in tumors from *Rosa26-CreER*$^{T2}$*; CiMKi*$^{WT/WT}$*, Rosa26-CreER*$^{T2}$*; CiMKi*$^{WT/TA}$*,* and *Rosa26-CreER*$^{T2}$*; CiMKi*$^{WT/KD}$ mice. **b)** Mutant expression in healthy skin from mice of all *CiMKi* genotypes, one, two, three, and four weeks after induction. See also Fig. 3c. **c-d)** tumour development over time (left graphs) and tumor incidence at end point (right graphs) of *CiMKi;Rosa26-CreER*$^{T2}$*;p53*$^{+/+}$ (c) or *CiMKi;Rosa26-CreER*$^{T2}$*;p53*$^{flox/+}$ (d).

**S4.**

**a)** Principal Component Analysis (PCA) plot showing the variance in gene expression profiles among all the samples. Each point represents a sample, and clustering indicates transcriptomic similarity across experimental groups. Note that some samples were sequenced using paired-end reads and others with single-end reads; both were included in the analysis following standardized normalization. 3 biological replicates were used in each condition, except for the high (TA/KD) CIN case at both timepoint.

**b)** Volcano plot displaying differentially expressed genes (DEGs) between high (TA/KD) CIN and no CIN skin at day 7 from CIN induction. Red dots represent significantly upregulated genes, and blue dots represent significantly downregulated genes (adjusted $p$-value < 0.05, $|\log_2$ fold change$| > 1$). Genes enriched in GO analyses are annotated.

**c)** Bar plot showing differentially expressed genes (DEGs) between High (TA/KD) CIN and no CIN skin on day 7 using Metascape enrichment tool. GO upregulated pathways in High CIN are related to keratinization and cell cycle, while fatty acids metabolism and cytokine expression are downregulated.

**d)** Heatmap (red high, blue down) showing expression of hair-follicle and epidermis specific keratin markers and fatty acids related genes at day 7 from CIN induction. Hair follicle keratins are upregulated in high (TA/KD) CIN, indicating hair follicle extension. For normalization, gene expression values were first averaged across the triplicates in the control group. All individual expression values (from both control and CIN levels) were then divided by this control average. The normalized triplicates values were averaged per group, and a $\log_2$ transformation was applied prior to visualization. Expression values are shown as relative fold changes compared to the control baseline.

**e)** Heatmap (red high, blue down) showing expression of hair-follicle and epidermis specific keratin markers and leukocyte activation genes at day 28 from CIN induction. Hair follicle keratins are upregulated in high (TA/KD) CIN, indicating hair follicle extension. Leukocyte activation genes are downregulated in high (TA/KD) CIN.

**f)** Bar plot showing differentially expressed genes (DEGs) between high (TA/KD) CIN and no CIN skin on day 28 using Metascape enrichment tool. GO upregulated pathways in high (TA/KD) CIN are related to cell cycle, while inflammation and immune system are downregulated.

**g)** Increased expression of Gli1 regenerative marker in the hair follicle cells (outer root sheath) in moderate (TA/TA) and high (TA/KD) CIN samples compared to no CIN skin after 28 days from CIN induction. Scale bars are 100 μm

**h)** Decrease of immune (CD45+) cells in moderate (TA/TA) and high (TA/KD) CIN skin compared to no CIN after 28 days from CIN induction. Scale bars are 100 μm.

**i-n)** Stainings as in figure 6, in order: Masson's trichrome, PCNAm ki67, SMA, CD45 and Cleaved-caspase 3. Scale bars are 100 μm.

**S5.**

**a)** Left: pictures of an example of a regressing and relapsing tumor in a *CiMKi*$^{KD/KD}$; *Rosa26-CreER*$^{T2}$ after 4-OHT treatment: regression is instantly, and almost complete at 18 days. Relapse is apparent 22 days after 4-OHT. Right: single curve of the tumor volume plotted against time for the example on the left. **b)** Curves of tumor volumes plotted against time for all single tumors (WT/WT (n=8), TA/TA (n=10), TA/KD (n=7), (KD/KD (n=19). For KD/KD: 2 out of 19 tumors that did nor regress are in black. **c)** *Mps1* mutant expression in two tumors from the same TA/KD mouse. Tumor 1 only expresses T649A (TA), and Tumor 2 only expresses D637A (KD).

**Table S1: PCR and Sequence primers**

| Gene | Forward primer | Reverse primer | Expected band size | Sequence primer | Mutation sequence |
|---|---|---|---|---|---|
| *CiMKi* (mutation) | GTGTCCTCACCC TGAAAATG | CAAAGCACAGCT GGGCTGTAGAG | ~1000 bp | CGGATTTTATTT TGAAGGTATTG | T649A (ACAàGCA ) D637A (GATà GCT) |
| *CiMKi* cDNA | CCTAGAAGACGC CGATAGCC | GTCTCTGATTGC TTCTGGGGC | ~400 bp | GATAAGATCAT CCGCCTCTATG | T649A (ACAàGCA ) D637A (GATà GCT) |

| | | | | | |
|---|---|---|---|---|---|
| Rosa26-CreER$^{T2}$ (mutant) | GGCAGGAAGCACTTGCTCTCCC | CCTGATCCTGGCAATTTCG | ~825 bp | NA | NA |
| Rosa26-CreER$^{T2}$ (wild-type) | GGCAGGAAGCACTTGCTCTCCC | GGAGCGGGAGAAATGGATATG | ~650 bp | NA | NA |
